# Problem-Solving as a Language: A Computational Lens into Human and Monkey Intelligence

**DOI:** 10.1101/2024.04.12.589234

**Authors:** Qianli Yang, Zhihua Zhu, Ruoguang Si, Yunwei Li, Jiaxiang Zhang, Tianming Yang

**Affiliations:** Institute of Neuroscience, Key Laboratory of Brain Cognition and Brain-inspired Intelligence Technology, Center for Excellence in Brain Science and Intelligence Technology, Chinese Academy of Sciences, Shanghai, China; Cardiff University Brain Research Imaging Centre, School of Psychology, Cardiff University, Maindy Road, Cardiff, CF24 4HQ, UK; University of Chinese Academy of Sciences, Beijing, China; School of Mathematics and Computer Science, Swansea University, Swansea, SA1 8DD, UK

## Abstract

Human intelligence is characterized by our remarkable ability to solve complex problems. This involves planning a sequence of actions that leads us from an initial state to a desired goal state. Quantifying and comparing problem-solving capabilities across species and finding its evolutional roots is a fundamental challenge in cognitive science, and is critical for understanding how the brain carries out this intricate process. In this study, we introduce the Language of Problem-Solving (LoPS) model as a novel quantitative framework that investigates the structure of problem-solving behavior through a language model. We adapted the classic Pac-Man game as a cross-species behavioral paradigm to test both humans and macaque monkeys. Using the LoPS model, we extracted the latent structure — or grammar — embedded in the agents’ gameplay, revealing the non-Markovian temporal structure of their problem-solving behavior. The LoPS model captured fine-grained individual differences among the players and revealed the striking differences in the complexity and hierarchical organization of problem-solving behavior between humans and monkeys, reflecting the distinct cognitive capabilities of each species. Furthermore, both humans and monkeys evolved their LoPS grammars during learning, progressing from simpler to more complex ones, suggesting that the language of problem-solving is not fixed, but rather evolves to support more sophisticated and efficient problem-solving. Through the lens of a language model, our study provides insights into how humans and monkeys break down problem-solving into compositional units and navigate complex tasks. This framework deepens our understanding of human intelligence and its evolution, and establishes a foundation for future investigations of the neural mechanisms of problem-solving.

## 2 Introduction

Advanced problem-solving is a hallmark of human intelligence, enabling us to navigate complex tasks to reap greater rewards and evade risks more effectively [1]. It has its roots in our primate ancestry and is likely shared by living nonhuman primates to varying degrees. Although many studies have investigated problem-solving capabilities in humans [2–5], quantifying and comparing problem-solving capability and its learning process across species remains a challenge. This is critical for us to understand the evolution of human intelligence and the underlying neural circuitry.

Problem-solving can be conceptualized as a process in which one establishes a sequence of operators or actions linking an initial state and a desired goal state. We hypothesize that this process, at its core, involves a systematic and structured process akin to a language. This language encapsulates the rules and principles that govern the composition and abstraction laws that construct the sequence [6–8] and guide our problem-solving efforts. The language may evolve to be more complex to support more sophisticated and more efficient problem-solving, both during evolution at a species level and during learning at an individual level.

To study this language of problem-solving quantitatively, we face two challenges. First, appropriate cross-species behavioral paradigms are needed for eliciting problem-solving behavior with rich structures [9–11]. Second, one has to develop a method to extract the structure of problem-solving from the agents’ behavior, which ideally does not depend on the agents describing how they solve the problem. Subjective reports are known to be inconsistent with actual problem-solving processes [12, 13] and are impossible in animal studies.

To address these challenges, we adapted the classic Pac-Man game as a cross-species behavior paradigm to test both humans and macaque monkeys. We developed a framework named Language of Problem Solving (LoPS) and developed a grammar induction algorithm to extract the underlying grammars — the non-Markovian temporal structure [14] — from the agents’ gameplay. The complexity of the resultant structure offers a quantifiable and interpretable metric of players’ problem-solving capacity [15].

Our results reveal striking differences in the complexity and hierarchical organization of problem-solving behavior between humans and monkeys and among individuals. Human players, especially the expert players, exhibited complex LoPS grammars with deep hierarchies. The grammar complexity correlated with the players’ performance, with more complex grammars associated with better problem-solving abilities. The grammar complexity was further manifested by a larger and more interconnected task state space in the expert players. Furthermore, both humans and monkeys evolved their problem-solving capabilities during learning, reflected in their grammars progressing from simpler to more complex.

Through the lens of a language model, our study offers a structured and systematic approach to understanding the complex cognitive process behind problem-solving. It reveals how humans and monkeys break down decision-making into compositional units and provides a framework for understanding how they navigate problem-solving [16–18], deepening our understanding of human intelligence and its evolution. The quantitative nature of the study establishes a foundation for future investigations of the neural mechanism of problem-solving.

## 3 Results

### 3.1 Pac-Man task

We modified the classic Pac-Man game for a cross-species study involving humans and monkeys. The game is complex enough to elicit rich and varied strategies, yet its core concept – foraging, hunting, and escape – is straightforward for monkeys to understand. In this game, players guide Pac-Man in a maze to eat pellets while avoiding and, at times, hunting ghosts. Two ghosts are programmed into the game. If caught by a ghost, players receive a time-out penalty, after all characters are reset to their starting locations. However, Pac-Man may eat a special pellet called an energizer, to temporarily turn the ghosts into a scared mode, during which they can be eaten by Pac-Man for extra rewards. Each game begins with randomly placed pellets and energizers. Players complete a game by clearing all the pellets in the maze and receive additional rewards.

We tested humans and monkeys with comparable game settings (See Methods 5.2), with one notable difference — the monkeys received real-time juice rewards corresponding to their in-game points, while the humans were shown a real-time cumulative point of the current game tally on their screens. During the game, we recorded all the game states and the players’ actions. More details are available in the Methods and our previous work [19]. In total, we collected behavioral data from a total of 34 humans and two monkeys.

### 3.2 Language of Problem-Solving

To play the game, the subjects need to create a solution by piecing together a sequence of actions to clear the maze. Our goal is to investigate this sequence with the LoPS model and extract the sequence structure, or the grammar of the LoPS, based on the observed problem-solving behavior and employ this knowledge to understand how individuals solve the game.

To achieve this goal, we first need to determine at which level we should build the language model. Although the Pac-Man game is solved by a sequence of actions (eg, ‘↑ ← ↓ →’ clears a square maze counter-clockwise), the basic components of the LoPS shall be at a higher level of abstraction. As motor actions depend too much on many details of game states, such as maze types and Pac-Man’s exact location, cognitively similar solutions can be associated with a large variety of action sequences, which prevents us from understanding the compositional principles of players’ problem-solving.

Our previous work [19] provides a better alternative. In that study, we have demonstrated a set of heuristic strategies that can account for the monkeys’ behavior in the Pac-Man game. Each strategy focuses on a subset of the game elements and a subgoal, and dictates the joystick action selection accordingly.

For example, with the *local* strategy, a player only considers the reward within a 10-tile radius and moves the joystick to guide Pac-Man in the direction that offers the largest reward, while the *evade* strategy directs Pac-Man away from the nearest ghost. Importantly, we showed that the monkeys’ gameplay could be described with sequences of dominating strategies.

We repeated the same analyses for the human players and found that the human players’ gameplay can be similarly described with sequences of dominating strategies. In addition, we identified two additional strategies: *stay* (keeping Pac-Man near a specific location) and *save* (keeping Pac-Man away from the energizer). These strategies allow players to take advantage of energizers and hunt ghosts efficiently.

With these seven strategies (Supplementary Table C4), we were able to accurately fit the joystick actions of both human and monkey players (Human: 90.6±3.0%, Monkey: 93.9±2.2%). Similar to the monkeys, the human players also employed Take-The-Best heuristics, dynamically choosing a single strategy at any moment (Supplementary Figure A1). Notably, the strategy fitting results indicate that two strategies, *stay* and *save*, were exclusively used by human players but not by monkeys.

Thereby, we converted both humans’ and monkeys’ gameplay into sequences of dominating strategies, upon which we developed the LoPS framework. The ‘words’ in the LoPS are a set of basic decision-making schemes (i.e. strategies), while the ‘grammars’ dictate how these strategies are concatenated into problem-solving sequences (Figure 1a).

**Figure 1.**
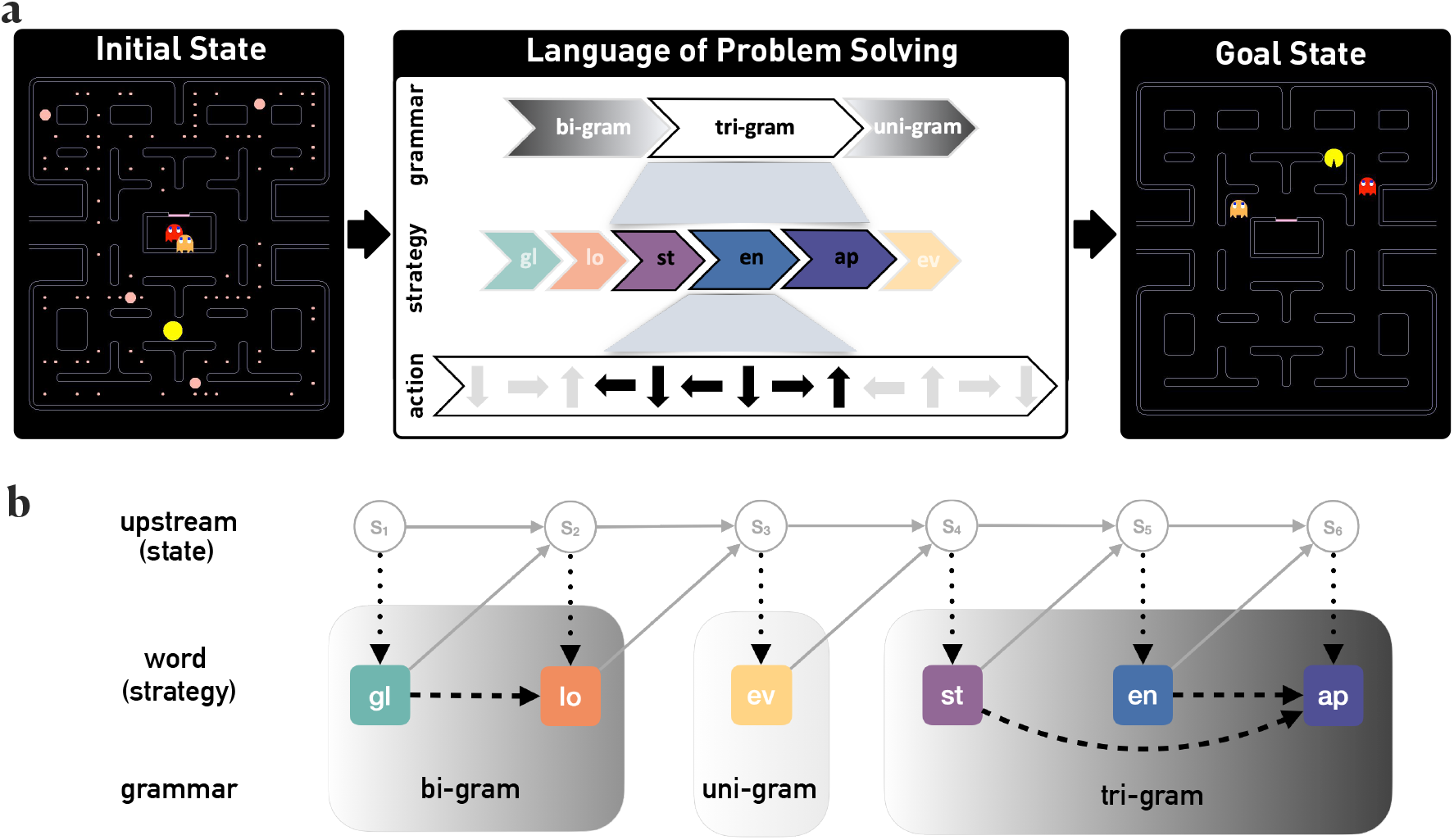
Language of Problem-Solving model. (a) The LoPS model conceptualizes problem-solving into several hierarchies of sequences that transform an initial state into a desired goal state. At the top hierarchy, a sequence of grammars, which describes game plans and governs how words (strategies) are concatenated. At the bottom level, strategies are implemented by actual actions of joystick movements. The strategy abbreviations are lo: *local*, gl: *global*, ev: *evade*, st: *stay*, ap: *approach*, en: *energizer*. Detailed definition of these strategies can be found in Supplementary Table C4, Supplementary Material A.2.1, and our previous work [19]. (b) UpN-gram. An upstream of game states evolves according to the game rules and the strategies applied (grey arrows). As a probabilistic graphical model, where nodes represent variables and edges denote the temporal dependency between them, a UpN-gram describes the history dependency of the words in addition to the upstream variables. Dotted edges denote the strong dependency between strategies and game states. Dashed edges denote the additional temporal dependencies, the grammars, considered by the UpN-gram.

### 3.3 UpN-gram

The core of the LoPS framework is a language model termed UpN-gram, which is based on the n-gram language model [20]. Like in a typical n-gram model, the probability of adopting a strategy in UpN-gram depends on its preceding *n*−1 strategies. In a uni-gram model (*n* = 1), each strategy is history-independent. With a bi-gram model (*n* = 2), strategies are Markovian, dependent only on the immediately preceding strategy. For more complex n-grams where *n >* 2, strategies are no longer Markovian, exhibiting dependencies on multiple preceding strategies. We also consider a variant, the skip-gram, in which strategies can depend on non-consecutive preceding strategies, skipping one or more in between. Long n-grams and skip-grams provide more flexible and efficient decision-making and demand more cognitive capacity.

Yet, unlike in standard n-grams, the words in the LoPS, the strategies, also depend on an upstream variable, the game states, which by themselves have a temporal structure. Because strategies can be in theory completely determined by the current game state, ignoring the temporal structure in the game states lead to false identification of strategy structures. Therefore, the dependency between strategies considered in our UpN-gram model has to be in addition to the dependency between strategies and states. That is to say, we are asking whether the knowledge of previous strategies provides further information on the current strategy in addition to what we know about the current game state. UpN-gram takes into account the game states as the upstream variable (Figure 1b). A uni-gram in UpN-gram means that the current strategy can be completely determined by the current game state, while a bi-gram indicates that the choice of the current strategy depends on not only the current game state but also the choice of the preceding strategy (Figure 1b). We recursively computed all concatenations of strategies with a significant dependency and induced the UpN-grams as the LoPS grammars of the agents (See 5.4 in Methods and algorithm 1). When applied to the synthetic data, our grammar induction algorithm can successfully recover the grammar used to generate the data (See A.2.4 in Supplementary Material).

### 3.4 LoPS grammars reveal different levels of complexity in players’ behavior

The induced LoPS grammar provides a window into the problem-solving structure of individual players. We first examined the LoPS ‘grammar books’ of the two monkeys. In order to visualize them, we organized the induced grammars hierarchically, ranked vertically based on their length (Figure 2a). Both monkeys exhibited simple grammars primarily consisting of uni-grams and bi-grams. For example, the *local* -*global* and *global* -*local* bi-grams suggest a divide-and-conquer strategy, where the monkeys first clear a local patch of pellets and then move to other pellet-rich regions to complete a game. The *energizer* -*approach* bi-gram indicates that the monkeys planned ahead to hunt the ghosts before consuming an energizer. The *evade* strategy is not used in any combinations, suggesting its impromptu nature. Other than that, the monkeys’ grammars can be divided into two categories: pellet-collection grammars and ghost-hunting grammars, each addressing a distinct aspect of the game.

**Figure 2.**
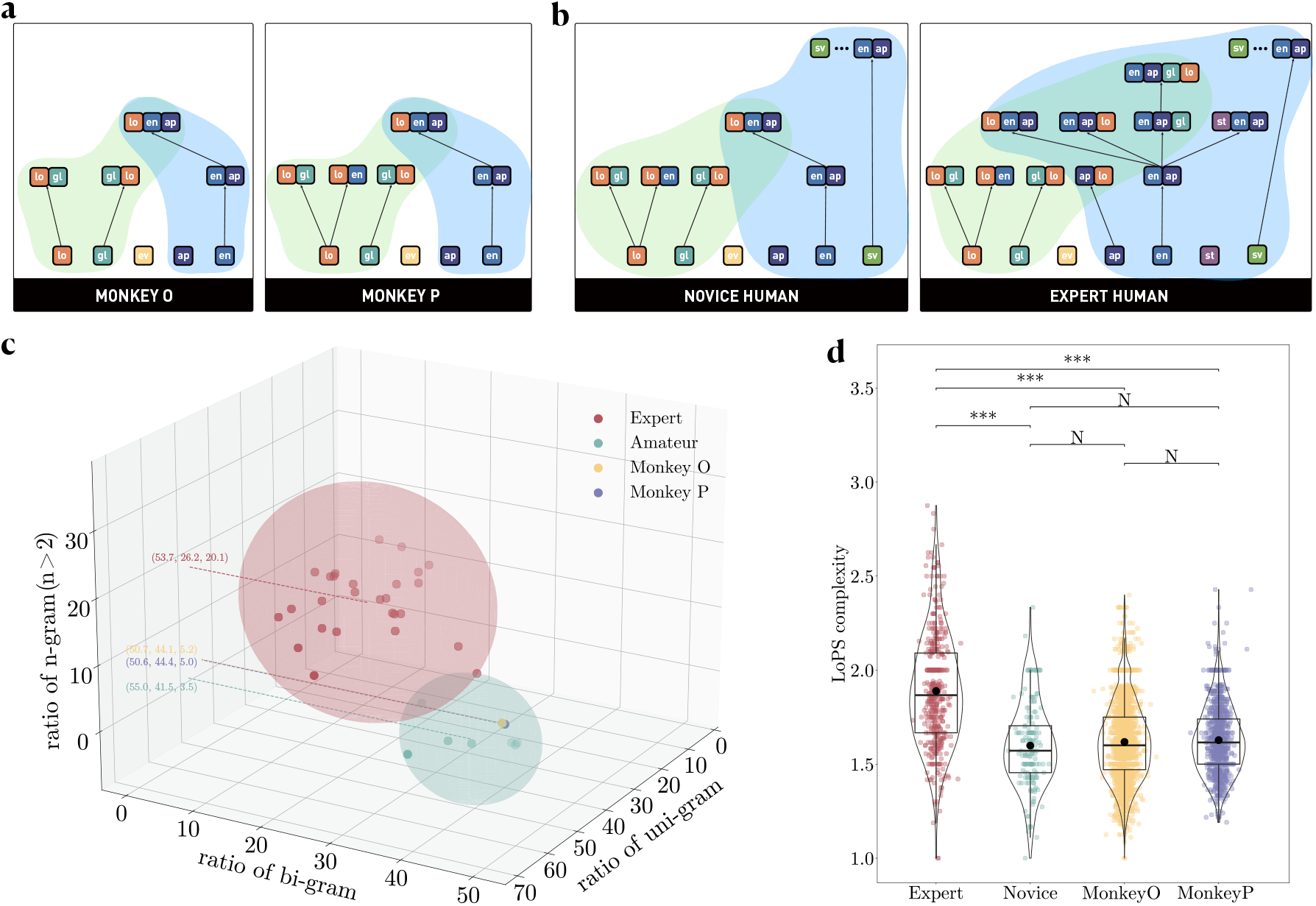
Participants’ LoPS grammars. (a) The LoPS grammars extracted from the two monkeys. Left: monkey O; right: monkey P. Each block is a basis strategy, i.e. uni-gram (lo: *local*, gl: *global*, ev: *evade*, ap: *approach*, en: *energizer*). Building from uni-grams, each high-level grammar is composed of two sub-grams. For clarity, only one line is shown, connecting to either the longer sub-gram or the first sub-gram if the two component sub-grams are of the same length. The green and the blue shading indicate the pellet-collecting and the ghost-hunting grammars, respectively, and the overlapping region indicates grammars combining pellet collection and ghost-hunting. (b) The LoPS grammars of two example human players. Left: example player 1, a novice; right: example player 2, an expert. Same convention as (a). Two additional basis strategies were used by the human players (sv: *save*, st: *stay*). (c) N-gram usage ratio. Each subject’s uni-gram, bi-gram, and complex n-gram (*n >* 2) usage ratio is plotted in a 3-D space. By applying the Agglomerative clustering method to n-gram usage ratios, we identified two distinct clusters among the human subjects: experts (red, *N* = 27) and novices (green, *N* = 7). The monkeys are indicated by the yellow (monkey O) and the purple dot (monkey P). Example gameplay videos can be found in Supplementary Videos 1-3. (d) LoPS complexity for experts, novices, and monkeys. Each data point represents the average LoPS complexity in a completed Pac-Man game. *** *p <* 0.001, NS *p >* 0.05 (Bonferroni-adjusted Mann-Whitney U tests).

Compared to the monkeys, human players exhibited more diverse LoPS grammars (Figure 2). Example player 1 added a *save* strategy, which leads to the skip-gram *save*…*energizer-approach*. This skip-gram helps the player to better utilize the energizer. Yet, example player 1’s LoPS grammars were overall similar to those of the monkeys (Figure 2b, left). In contrast, example player 2 developed much more sophisticated grammars, including a large number of tri-grams and even a quad-gram (Figure 2b, right). Many of these grammars were specifically for ghost-hunting. For instance, the *stay-energizer-approach* tri-gram indicates that the player lured ghosts while staying near an energizer, then ate the energizer and swiftly hunted the ghosts (Supplementary Video 5). In addition, a large number of long grammars, such as *energizer-approach-local-global*, string together pellet-collection and ghost-hunting grammars, demonstrating a high level of planning that combines the two most important aspects of the game.

The two example human players illustrate the marked diversity among human players’ LoPS grammars, ranging from simple monkey-like grammars to highly sophisticated ones. In Figure 2c, we plot the uni-gram, bi-gram, and tri-gram usage ratios for all human players in a 3-D space. Agglomerative clustering reveals two clusters. Example human player 1 belongs to the first cluster, which favored simpler uni-grams and bi-grams (Uni-gram: 55.0±3.8%, bi-gram: 41.5± 4.5%, n-gram (*n >* 2): 3.5± 1.9%), while the participants in the second cluster, including example human player 2, demonstrated a much higher usage ratio for tri-grams (Uni-gram: 53.7 ±5.5%, bi-gram: 26.2 ±5.2%, n-gram (*n >* 2): 20.1 ±4.0%). We refer to the players in the two clusters as novice and expert players, respectively.

Interestingly, the two monkeys, when plotted in the same 3-D space (monkey O, uni-gram: 50.7±3.7%, bi-gram: 44.1 ±3.5%, n-gram (*n >* 2): 5.2± 1.1%; monkey P, uni-gram: 50.6 ±6.5%, bi-gram: 44.4 ±6.1%, n-gram (*n >* 2): 5.0± 1.5%), fall into the cluster of novice players, suggesting that they share similar problem-solving structures. Notably, however, both monkeys had been playing the game for three years, while some of the novices played the game for the first time in their life. Exemplar gameplay videos from different groups of humans and monkeys can be found in Supplementary Videos 1-3.

To further quantify the players’ grammar complexity, we developed a summary statistic, which we term LoPS complexity. It is defined as the average of the grammar length, 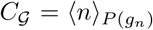, calculated over the parsed n-gram sequence 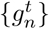 based on each player’s grammar structure. Figure 2d plots the LoPS complexity for the two human groups and the monkeys. As expected, the expert humans, the novice humans and the two monkeys showed significant differences in LoPS complexity (*H*(3, 4647) = 155.30, *p <* 0.001). Further pair-wised contrast suggests that the expert humans showed significantly higher average LoPS complexity than both the novice humans and the monkeys (all *p <* 0.001, Bonferroni corrected).

### 3.5 Complex grammar correlates with better game performance

Complex grammars come at a computational cost, and their existence can only be justified if they provide behavioral advantages. Indeed, we found a strong positive correlation between LoPS grammar complexity and problem-solving performance at the individual level (Figure 3a; Pearson’s *r* = 0.65, bootstrap 95% CI (0.45, 0.82)). With the help of more complex grammars, the expert humans significantly outperformed the novice humans (*U* = 0.00, *p <* 0.001, Mann-Whitney U test). The two monkeys, when plotted on the same graph, fall on the lower end of both grammar complexity and performance, suggesting a continuum between humans’ and monkeys’ problem-solving abilities.

**Figure 3.**
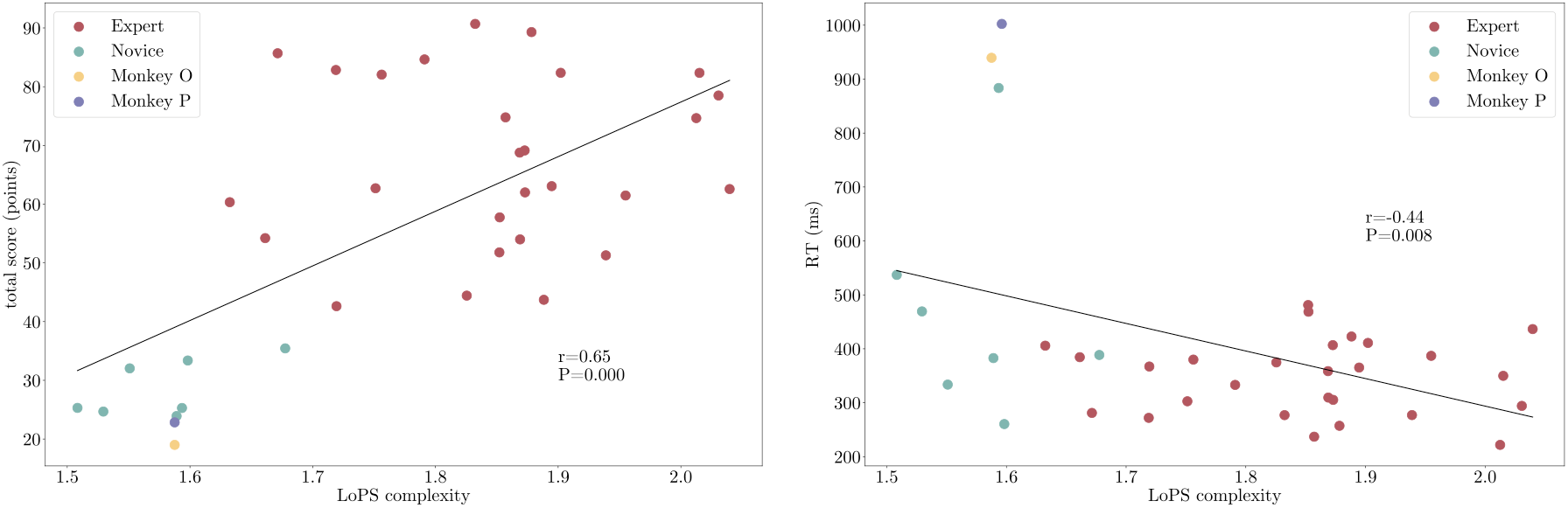
Grammar complexity correlates with performance. (a) Performance score. A significant positive correlation is observed between LoPS complexity and total score (Pearson’s *r* = 0.65, bootstrap 95% CI (0.45, 0.82), excluding the monkeys). The total score is calculated as the accumulated points gained by completing one game, excluding points from pellets and only counting those related to energizers, ghosts, and death (detailed in Supplementary Table C1). Each data point represents one subject. (b) Reaction time. A significant negative correlation is observed between LoPS complexity and reaction time (Pearson’s *r* =−0.44, bootstrap 95% CI (−0.75, − 0.25), excluding the monkeys). Reaction time (RT) is defined as the time between the occurrence of a specific event (such as energizer or ghost consumption) and the subject’s first Pac-Man direction change. The lines are from linear regressions using the human data. Red: expert players, green: novice players, yellow: monkey O, purple: monkey P.

In addition to game scores, we also examined how fast an agent may respond to certain game events. While it is difficult to measure the reaction times of strategy switches in general, we observed that the agents often changed Pac-Man’s direction immediately after eating an energizer or a ghost, indicating a switch of strategy. We measured the onset of these direction changes as the reaction time and examined if they correlated with grammar complexity. We hypothesized that agents with complex LoPS grammars could offload the real-time decision-making by planning strategy switches well in advance, leading to faster reaction times. Consistent with this hypothesis, we found a significant negative correlation between LoPS complexity and reaction time (Figure 3b; Pearson’s *r* =− 0.44, bootstrap 95% CI (−0.75, − 0.25)). The Kruskal-Wallis test on two human groups and two monkeys indicated a significant difference between these four groups (*H*(3, 6212) = 517.62, *p <* 0.001). Further pair-wised comparison suggested that the expert humans exhibited shorter reaction times than the novices and both monkeys (all *p <* 0.001, Bonferroni corrected).

These results suggest that complex LoPS grammars provide both performance and efficiency benefits in problem-solving.

### 3.6 LoPS grammar evolves with learning

Both monkeys and humans became better at the game with experience. Employing the LoPS induction algorithm at different stages of the experiment, we can answer the question whether their performance improvements were reflected in their LoPS grammars.

The monkeys were tested continuously over a span of three years. Through years of gaming, the monkeys continued building their grammar set and adding longer grammars (Figure 4ab). The new grammars, such as the *energizer-approach* bi-gram and the *local-energizer-approach* tri-gram, were primarily related to ghost-hunting. They helped the monkeys take better advantage of energizers and made their behavior more human-like.

**Figure 4.**
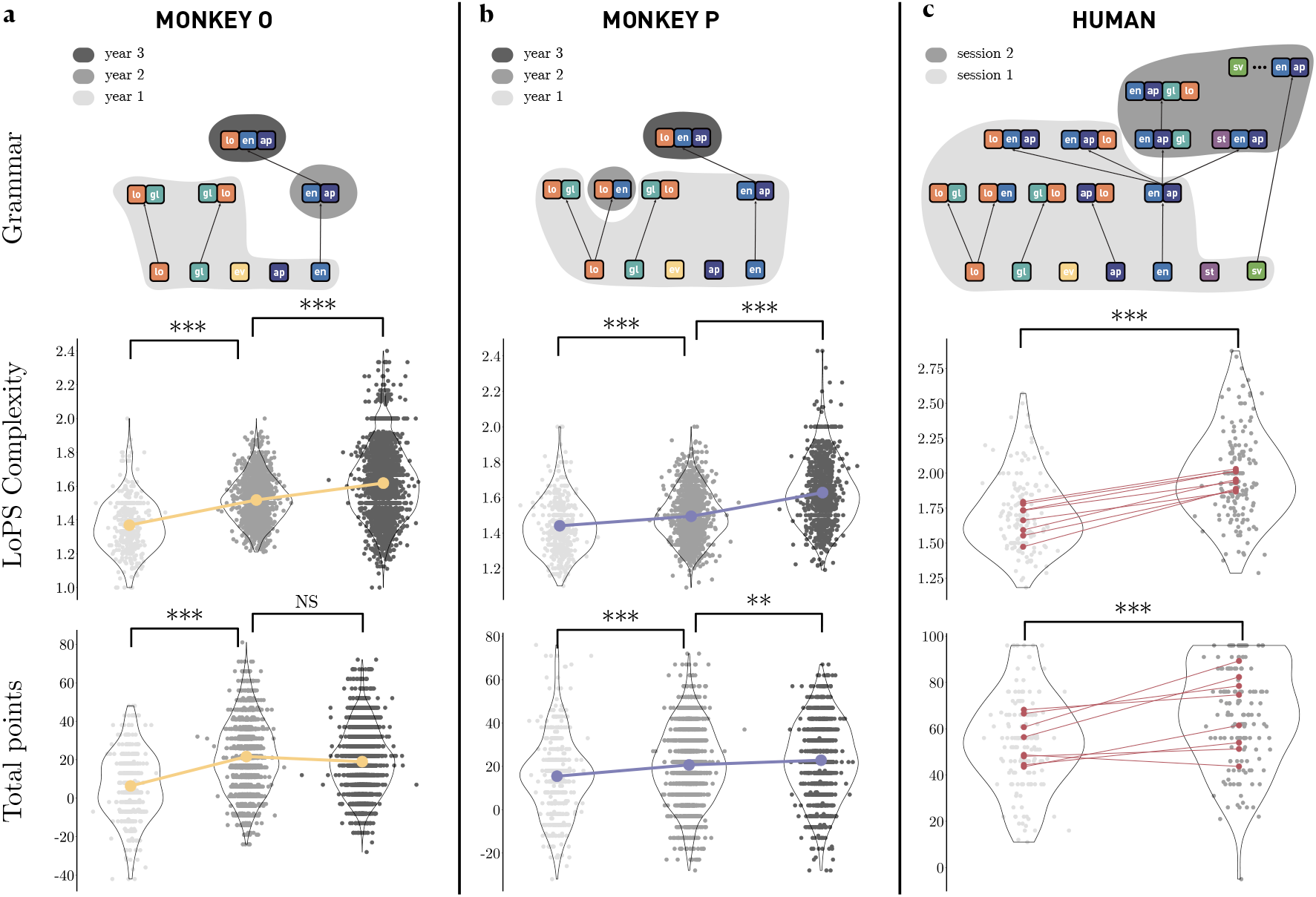
Evolving LoPS grammars with learning. (a) Monkey O. Top: LoPS grammars extracted from the monkey O’s behavior. The different gray shades indicate the additions of new grammars during each year. Middle: Progression of grammar complexity for each year. Bottom: Progression of game scores for each year. Each data point represents the mean in a complete Pac-Man game trial. The yellow dots represent the means of each year. (b) Monkey P. Same convention as in (a). (c) Human participants. We assessed LoPS grammar for all 34 human subjects over two sessions. The LoPS complexity for 8 of these subjects increased significantly in the second session (*p <* 0.001, two-sample t-test). Top: LoPS grammars of the 8 subjects with significant LoPS complexity increases. The different gray shades indicate the two sessions. Middle: Progression of grammar complexity in the two sessions. Bottom: Progression of game points in the two sessions. Each data point represents the mean in a complete Pac-Man game trial. The red dots are the means of all trials of the 8 subjects with significant LoPS complexity increases. *** *p <* 0.001, ** *p <* 0.01, NS *p >* 0.05 (two-sample t-test).

The human participants were tested in two sessions. We examined the evolution of their LoPS grammar across these sessions (Figure 4c). In 8 out of the 34 subjects, there was a significant increase in the complexity of their LoPS grammars between the sessions (Figure 4c, middle, *p <* 0.001, two-sample t-test). This increase in complexity was attributed to the emergence of more complex grams in the later session, rather than an increased frequency of grams already present in the first session (Figure 4c, top). Correspondingly, the task performance of these subjects improved significantly across the sessions (Figure 4c, bottom, *p <* 0.001, two-sample t-test). No subjects exhibited a decrease in their grammar complexity or task performance.

These results unveiled the expansion of the LoPS grammar set in both the monkeys and humans during learning, indicating their problem-solving capacities evolved with experience. Yet, new strategies emerged in the monkeys only after years of practicing, and the monkeys’ LoPS grammars remained much less sophisticated than those of the expert humans. There appears to be a performance ceiling separating monkeys and humans, reflected in the complexity of their LoPS grammar.

### 3.7 Structure of state variable dynamics

LoPS grammars describe sequences of strategies, which directly lead to changes in game states. Therefore, LoPS not only captures the structure of strategy sequences but also should account for how the game states change as strategies are executed. To investigate the relationship between LoPS grammars and game state structure, we examined how different state variables co-varied after the execution of a strategy in the experts, novices, and monkeys — three groups of players with distinct grammar patterns. We focused on six state variables that are most essential to the game (See definitions in Supplementary Table C3) and computed their joint probabilities, denoted as *P* (s). Using the PC algorithm [21, 22], we induced the Markov network that depicted the correlational structure between these state variables (See Method 5.6). The resulting graph describes the inter-dependency between these state variables as the result of the execution of strategies.

We observed distinct structures for expert humans, novice humans, and monkeys (Figure 5). The latter two groups formed two separate state variable clusters (Figure 5, middle and right). One cluster consisted of two interdependent state variables, the distance to the energizer and the count of local pellets. The other cluster consisted of four ghost-related variables, each ghost’s mode and Pac-Man’s distance to each ghost. What is interesting is the missing link between the two clusters in the novices and monkeys. In the experts, the two clusters were linked together via *e*, the distance-to-energizer (Figure 5, left). Thus, the experts created a graph structure with the energizer as the bottleneck, connecting the ghost-related and the foraging-related state variables (Figure 5). This is reflected by their large number of long grammars that combine pellet-collection and ghost-hunting strategies. They are absent in both the novices and the monkeys.

**Figure 5.**
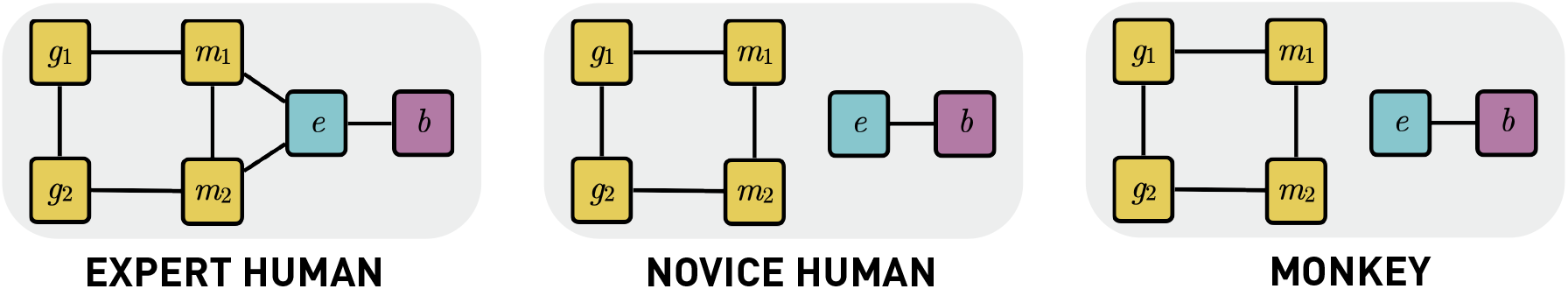
Markov networks of game state variables in the experts (left), novices (mid), monkeys (right). The nodes represent the six state variables: distances from Pac-Man to each ghost (*g*_1_, *g*_2_), modes of the ghosts (*m*_1_, *m*_2_), Pac-Man’s distance to the energizer (*e*), and counts of local pellets (*b*). Edges between the nodes indicate conditional dependencies. The PC algorithm [21, 22] was used to induce the graph structure (Supplementary Material A.4).

### 3.8 State space

To further investigate how LoPS grammars capture the game state dynamics, we examined the transitions of game states. Game states are defined as the ensemble of the state variables. Again, we focused on the six essential state variables. For each group of players, we identified all the states that appear in their gameplay with a probability above the random chance threshold and plotted the states in a circular configuration (Figure 6). The states from the expert humans’ gameplay greatly outnumber those from the novices and the monkeys’ (experts: 19, novices: 7, monkeys: 5), leading to a much more complex transition map. Many of the states that are unique to the experts take energizers into account (states 6-22), while the gameplay of novices and monkeys mainly consists of states that disregard the distance to the nearest energizers (states 1-5). This highlights the importance of energizers in the expert’s game. By adopting more sophisticated grammars, the expert players were able to access game states that were more advantageous but unexplored by the novice or monkey players. The ability to solve a problem in a larger state space is indicative of a greater problem-solving capacity.

**Figure 6.**
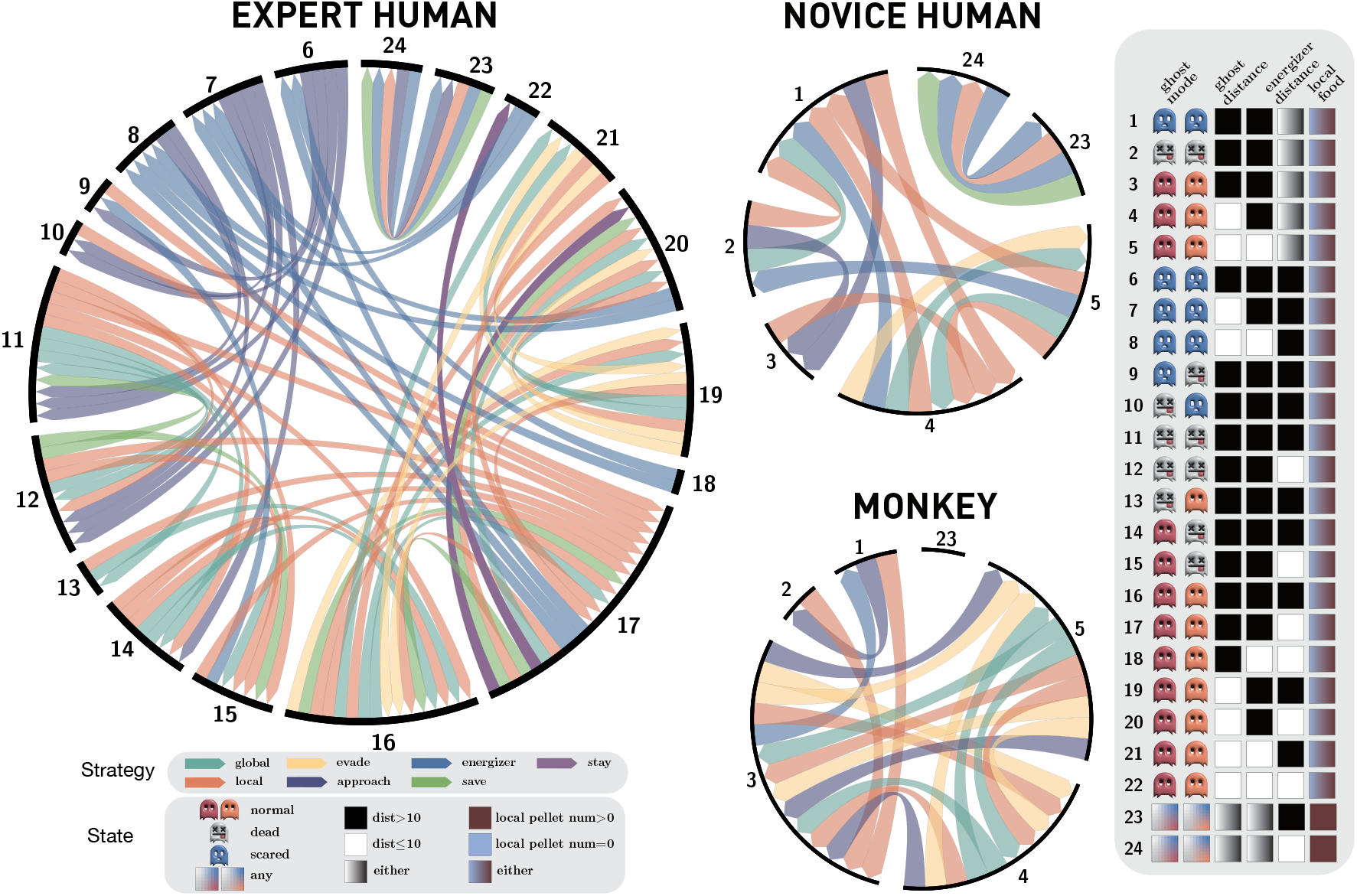
State spaces. Circular representations of game state spaces are plotted for the experts, novices, and monkeys. Game states are defined as vectors of six state variables (right inset). The distance and the pellet count variables are binarized for simplicity (left bottom inset). In addition, a state can have a special value of *any* for a particular variable, indicating that the state can take any value for that variable. Only states with a probability higher than the chance level are plotted (experts: 19 states, novices: 7 states, monkeys: 5 states). Directed edges denote strategies that lead to state transitions.

## 4 Discussion

Our investigation into the cognitive processes underpinning problem-solving behavior through a language model has yielded intriguing insights into the differences between human and nonhuman primate intelligence. The adaptation of a classic video game like Pac-Man into a cross-species behavior paradigm allows us to explore these processes in a dynamic and quantifiable manner. The model’s ability to capture the temporal structure and complexity of problem-solving brings us closer to understanding the compositional nature of cognition, both in humans and macaque monkeys.

### 4.1 Language of Thoughts

The LoPS model draws inspiration from the language of thoughts (LoT) hypothesis, which conceptualizes mental representations through the lens of symbolic primitives and composition laws [6]. This hypothesis has effectively captured core principles across a wide range of human cognitive abilities, including concept learning [23], sequence learning [15], and artificial handwriting generation [24]. The LoPS model offers an empirical bridge to the theoretical constructs of the LoT hypothesis, enriching the LoT literature with data-driven evidence and providing a concrete framework through which we can understand and analyze the ‘syntax’ and ‘grammar’ of cognitive processes involved in problem-solving. It offers a new perspective on how compositional thought processes can be measured and compared across different species.

Furthermore, the differences in LoPS grammar complexity between humans and monkeys provide empirical support for species-specific ‘languages of thought’ [25]. Our study suggests that while there may be a shared ancestral cognitive language, the depth and complexity of these mental languages have evolved significantly, leading to the sophisticated problem-solving abilities observed in humans.

### 4.2 Human vs Monkey

Although the intelligence gap between humans and other animals, including monkeys, is undisputed, the extent and underlying causes of this separation are still a subject of ongoing research [26]. Our results clearly demonstrate that human players exhibit more complex problem-solving behaviors compared to monkeys, and the hierarchical nature of human cognition [27–30] is mirrored in the LoPS grammars, which are more intricate and possessed deeper hierarchies. These findings align with the notion that human intelligence is distinguished by its ability to use abstract thought processes and to construct and navigate complex state spaces capabilities that are less developed in macaque monkeys. Moreover, humans displayed a capacity for quickly developing new complex grammars in their tasks through experience, which further increased their performance advantage over monkeys. Together, our results suggest that hierarchical organization in cognitive processes is a significant factor in the evolutionary advantage of human intelligence.

Interestingly, both humans and monkeys demonstrated an evolution in their problem-solving strategies, shifting from simpler to more complex grammars as they learned. This supports the idea that intelligent behavior is not static; it evolves and adapts in response to experience and learning. It also reinforces the concept that problem-solving is not a fixed ability but a dynamic skill that can be developed and refined over time.

### 4.3 UpN-gram

We established the UpN-gram language model to capture the influence of the underlying state transitions on the temporal structure [31]. This is the key to identifying the true temporal dependencies of strategies. Our grammar induction algorithm is a significant improvement over the previous methods that primarily focused only on action sequences and ignored the temporal structure of the internal and external variables that lead to the actions [32]. In most problem-solving scenarios, ignoring these variables and their temporal structures would lead to false identifications of solution structures.

While the UpN-gram model is simple and intuitive, it does have some limitations. One key constraint is its inflexibility as it only factors in history of a set length. This limits the model’s ability to capture other types of dependencies in the data. In addition, the n-gram model can suffer from data sparsity, as some n-grams might be rare in training datasets. One potential way to solve these problems is to incorporate other logical operations to diversify the composition rules over primitives. The And-Or graph grammar is a prominent alternative [33, 34], where the *and* operation captures conjunctions between elements, while the *or* function denotes choice alternatives. State-of-the-art language models based on the RNNs and transformers [35] present another promising avenue. These models accommodate a richer palette of primitives and more adaptable compositional rules [36].

### 4.4 Neural substrate of LoPS

The LoPS reveals a hierarchical structure in the problem-solving of humans and monkeys. A rostrocaudal axis of the PFC has been proposed to reflect the hierarchical cognitive control, with the brain implementing a deeply structured, tree-like policy by nesting multiple corticostriatal gating loops, arranged from back to front within the frontal lobe [37, 38]. Theories propose that this region coordinates the activation of lower-level regions based on higher-order cognitive representations [39]. Consistent with this idea, patients with lesions in the rostral PFC often exhibit impairments in problem-solving tasks that require planning and abstract reasoning [40]. In addition, the human brain’s language circuitry and its counterparts in the monkey brain may also play an important role in LoPS.

### 4.5 Conclusion

In conclusion, the LoPS framework provides a unique lens through which we can view and quantify problem-solving behavior. The distinctions between human and monkey problem-solving grammar underscore the advanced cognitive capabilities that characterize human intelligence. Our study not only contributes to our understanding of problem solving but also offers a framework for future explorations into the evolution of intelligence and the neural substrates of problem-solving.

## 5 Methods

### 5.1 Subjects and Materials

This study recruited 34 healthy participants (19 female, 14 male, and 1 undisclosed). The participants were aged from 18 to 40 with an average age of 24.79 ±5.45. None of the participants had a previous history of neurological or psychiatric disorders. All participants gave written informed consent for the sessions they attended and received monetary compensation. The study was approved by the Cardiff University School of Psychology Research Ethics Committee.

Two male rhesus monkeys (Macaca mulatta) were used in the study (O and P). They weighed on average 6-7 kg during the experiments. All procedures followed the protocol approved by the Animal Care Committee of Shanghai Institutes for Biological Sciences, Chinese Academy of Sciences (CEBSIT-2021004).

### 5.2 Pac-Man task

The Pac-Man game for the monkey experiment has been described previously [19]. Briefly, monkeys control Pac-Man with a joystick through a maze to collect pellets, each pellet consumed yields a juice reward. Completing the maze yields additional juice. Two ghosts roam the maze. If Pac-Man is caught, there is a time-out penalty. Eating special pellets, called energizers, switches ghosts into a temporary scare mode during which they can be eaten for extra juice rewards.

For the human task version, the human controls the Pac-man with a keyboard. The maze is sized 725×900 pixels and displayed at the resolution of 1920×1080. Same as the monkey task version, the maze is divided into square tiles of 25×25 pixels. The human game has an extra tile, which allows Pac-Man to start location to be located in the center of the maze horizontally. This modification does not change the connectivity of the maze. The human game also removes all the fruits to simplify the game. We define 9 unconnected pellet patches in the maze. Each patch has 9 to 13 tiles, and in each game, only 7 randomly selected patches are filled. Totally, there are 76 to 83 pellets in the maze, among which four randomly selected pellets at the upper-left, upper-right, lower-left, and lower-right areas of the maze are replaced with an energizer. Collecting a pellet and an energizer will gain 2 and 4 points of score, respectively. Moreover, humans will get an extra 10 points for catching a scared ghost. A real-time cumulative point of the current game tally is shown on the screens. If caught by a ghost, the humans receive a deduction of points. For better data collection, Humans are allowed maximally two attempts to complete a game. Finally, the game speed for the human version is set to be twice as fast as that of the monkey version. All the other settings are identical. The human-specific settings aim to encourage more evenly distributed strategies used by human subjects to facilitate the behavior analyses with limited amount of data.

The monkeys’ behavioral data was collected in three years. Across three years, the monkeys played 820 (Year 1), 3240 (Year 2), and 571 (Year 3) game trials. The analyses presented here are based on the behavior in Year 3 unless mentioned otherwise. On average, the monkeys completed 33±9 games per session, with each game requiring 4.9±1.8 attempts. The human participants were involved in two separate sessions. We used the behavior data in the second session to do the analyses in the paper unless mentioned otherwise. On average, each human participant completed 36 ± 6 games per session, with each game taking an average of 1.2 ± 0.4 attempts.

### 5.3 Language of Problem-Solving

The Language of Problem-Solving (LoPS) model is a language model that describes the structure of an agent’s problem-solving. Primitive words in the LoPS are a set of decision-making schemes, which are a selection of basis strategies in the context of the Pac-Man game. Grammars in the LoPS describe the temporal organization of strategy sequences.

In the current study, we established the UpN-gram based on the n-gram language model for the LoPS. In UpN-gram, the probability of adopting a strategy is dependent on its preceding *n*− 1 strategies in addition to the upstream variables, the game states. The UpN-gram can be mathematically formulated under the framework of Probabilistic Graphical Models (PGMs). A PGM *G* = {𝒱, *ℰ*}, contains nodes 𝒱 representing random variables, and edges E indicating conditional dependencies. The edges are either directed (𝒱_*k*_ → 𝒱_*l*_) or undirected (𝒱_*k*_ − 𝒱_*l*_), used in Bayesian networks and Markov networks, respectively.

An UpN-gram is defined as a Bayesian network 𝒢:

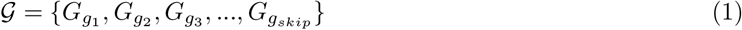

where 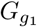 (uni-gram), 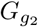 (bi-gram), 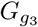 (tri-gram), …, and 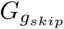 (skip-gram) can be defined as:

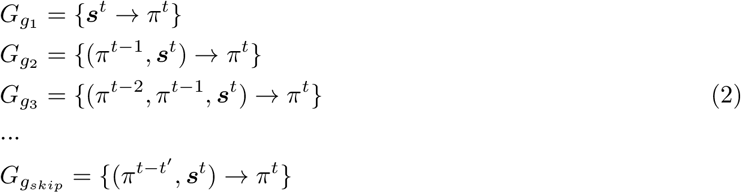

where s^*t*^ represents game state at time *t*, π^*t*^ represents strategy at time *t, t*^′^ *>* 1 denotes the temporal dependency of non-consecutive preceding strategies. All concepts used in the LoPS can be found in Supplementary Table C2.

### 5.4 LoPS gammar induction

Grammar induction is an inverse problem in which we commence with a subject’s gameplay and deduce the LoPS grammar. We first convert the action sequence into strategy sequence (Supplementary A.2.2) [19]. Together with the game states, we obtained the time series of strategy *D*_π_ and state *D*_*s*_. We then induce the grammar 𝒢. We start with a grammar graph that only contains uni-grams,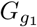. Then we compute the likelihood score of an alternative graph by concatenating one pair of strategies. If the likelihood score for the alternative graph is higher than that for the null graph, we accept the alternative graph. Upon each update of the grammar graph, we parse the strategy time series to generate a new UpN-gram time series, where the parsing algorithm follows the principle of Minimum Description Length (MDL). The new UpN-gram time series is used to compute the likelihood score in the subsequent iteration. This iterative process continues until the likelihood score for the grammar graph reaches the maximum. The induced graph structure is defined as the agent’s LoPS grammar 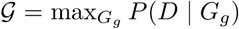. More details can be found in Supplementary Material A.2.

### 5.5 LoPS analyses

We create a subject’s grammar graph by organizing the induced grammars as the nodes hierarchically, ranked vertically based on their length (Figure 2a). Each higher-level grammar node is built from two simpler grammar nodes. For better visualization, only one link is shown. If a complex grammar is created from daughter nodes of equal complexity, only the edge to the first node is shown. For cases where a complex grammar breaks down into two daughter nodes of differing complexity, we retain the edge to the more complex grammar node. This approach provides a clear and concise visual representation, emphasizing the inherent hierarchical and compositional nature of the grammars.

Given each subject’s grammar book, we apply the parsing algorithm (Algorithm 5) on each individual’s sequence of strategies to obtain their respective n-gram sequences. Based on these n-gram sequences, we can compute the probability of each n-gram, denoted as *P* (*g*_*n*_). The ratio of all participants’ uni-gram, bi-gram, and tri-gram usage can be computed accordingly. Employing the agglomerative clustering method on these three statistical measures across 34 human subjects leads to the identification of two clusters, named experts and novices according to their grammar usage preference. The clustering algorithm used the Euclidean distance metric and “ward” linkage criterion to minimize the variance of the clusters being merged.

We define the complexity of each UpN-gram grammar *g*_*n*_ as its depth *n*. The complexity of a skip-gram corresponds to the length of the subsequent UpN-gram sequence up to the targeted grammar element. The LoPS complexity can then be computed as the average of *n* with respect to the probability of the n-grams, represented as 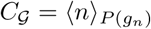. This measure provides a concise summary statistic to quantify the grammar complexity.

### 5.6 State variable inter-dependency

We use a Markov network to illustrate the correlational structure among game state variables (Figure 5). Nodes in the network graph are the six game state variables, denoted as s = (*s*_*m*1_, *s*_*m*2_, *s*_*g*1_, *s*_*g*2_, *s*_*e*_, *s*_*b*_), and undirected edges indicate conditional dependencies between nodes. Definition of these states can be found in Supplementary Table C3. To learn the graph structure, we use the PC algorithm (Algorithm 6, [21, 22]).

### 5.7 State space

Game state is defined as a vector of length six, combining the six state variables described above. The distance and the pellet-count variables are binarized for simplicity. In addition, a state can have a special value of *any* for a particular variable, indicating that the state can take any value for that variable.

Effective states are defined as state configurations with probabilities above the chance level *P* (*S*) *>* 1*/* ∣ *S* ∣.

We factorize the joint probability of state variables according to the structure of the state variable inter-dependency network. According to the experts’ network (Figure 5 left), the factorized joint probability is *P* (*s*) = *P* (*s*_*g*_|*s*_*e*_)*P* (*s*_*b*_|*s*_*e*_)*P* (*s*_*e*_), where 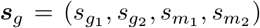. The state configurations are defined with respect to two tuples 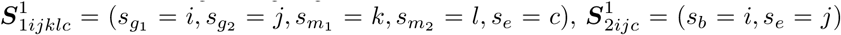, where *i, j, c* ∈ {0, 1} and *k, l* ∈ {0, 1, 2}. Thus in total, the potential state number is 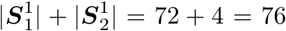.

For novices and monkeys (Figure 5 mid, right), the factorized joint probability is *P* (*s*) = *P* (*s*_*g*_)*P* (*s*_*b*_, *s*_*e*_). The state configurations are defined with respect to two tuples 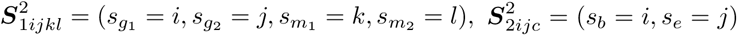, where *i, j* ∈ {0, 1} and *k, l* ∈ {0, 1, 2}. Thus in total, the potential state number is 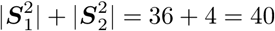.

A player’s game space is defined as the set of the player’s effective states. We then used the structure learning method (Supplementary Material A.3) to induce the transition graph among these effective states. For each transition {*S, S*^′^, π}, where *S*^′^ is the successor effective states to a preceding effective states *S*, with the subject employing the strategy π, we defined a null graph to be *G*_0_ = {*S, S*^′^, π} and alternative graph to be *G*_1_ = {(*S*, π) →*S*^′^}. In Figure 6, we illustrated all the transitions for which the likelihood score of the alternative graph exceeds that of the null graph, across each group of subjects.

## Acknowledgements

We thank Yunxian Bai, Wei Kong, Jianshu Li, Zhongqiao Lin, Ruixin Su, Lu Yu, Yang Xie, and Wenyi Zhang for their help in all phases of the study, and Thomas Akam for helpful discussions.

## Funding

This work was supported by National Science and Technology Innovation 2030 Major Program (Grant No. 2021ZD0203701) to T.Y., National Natural Science Foundation of China (Grant No. 32100832) to Q.Y., European Research Council (ERC) under the European Unions Horizon 2020 research and innovation programme (Grant agreement No. [716321 - FREEMIND]) to J.Z., and PhD studentship from China Scholarship Council to R.S.

The funders had no role in study design, data collection and interpretation, or the decision to submit the work for publication.

## Conflict of interest

The authors declare no competing financial interests.

## Data and code availability

The data and codes that support the findings of this study are provided at: https://github.com/superr90/LoPS.

## Author contribution

Qianli Yang: Conceptualization; Resources; Software; Formal analysis; Supervision; Funding acquisition; Validation; Investigation; Visualization; Methodology; Writing - original draft; Project administration; Writing - review and editing. Zhihua Zhu: Software; Formal analysis; Investigation; Methodology; Writing - review and editing. Ruoguang Si: Data curation; Investigation; Writing - review and editing. Yunwei Li: Data curation; Investigation; Writing - review and editing. Jiaxiang Zhang: Supervision; Funding acquisition; Writing - review and editing. Tianming Yang: Conceptualization; Resources; Software; Formal analysis; Supervision; Funding acquisition; Validation; Investigation; Visualization; Methodology; Writing - original draft; Project administration; Writing - review and editing.

## Appendix A Supplementary Material

### A.1 Take-The-Best strategy heuristic

Players could adopt one or a mixture of multiple strategies when playing the Pac-Man game. We assumed that the final decision for Pac-Man’s moving direction was based on a linear combination of the basis strategies and asked whether there was a dominating strategy. We estimated the strategy weights using time windows of flexible length, by assuming that the relative strategy weights are stable for a period. We ranked the strategies according to their fitted weights at each time segment. We observed that the weight difference between the first and the second most dominating strategies was heavily skewed toward one (Supplementary Figure A1). The result is consistent with our previous findings [19], indicating that both the humans and monkeys adopted Take-The-Best heuristics in which action decisions were formed with a single strategy heuristically and dynamically chosen.

**Supplementary Figure A1:**
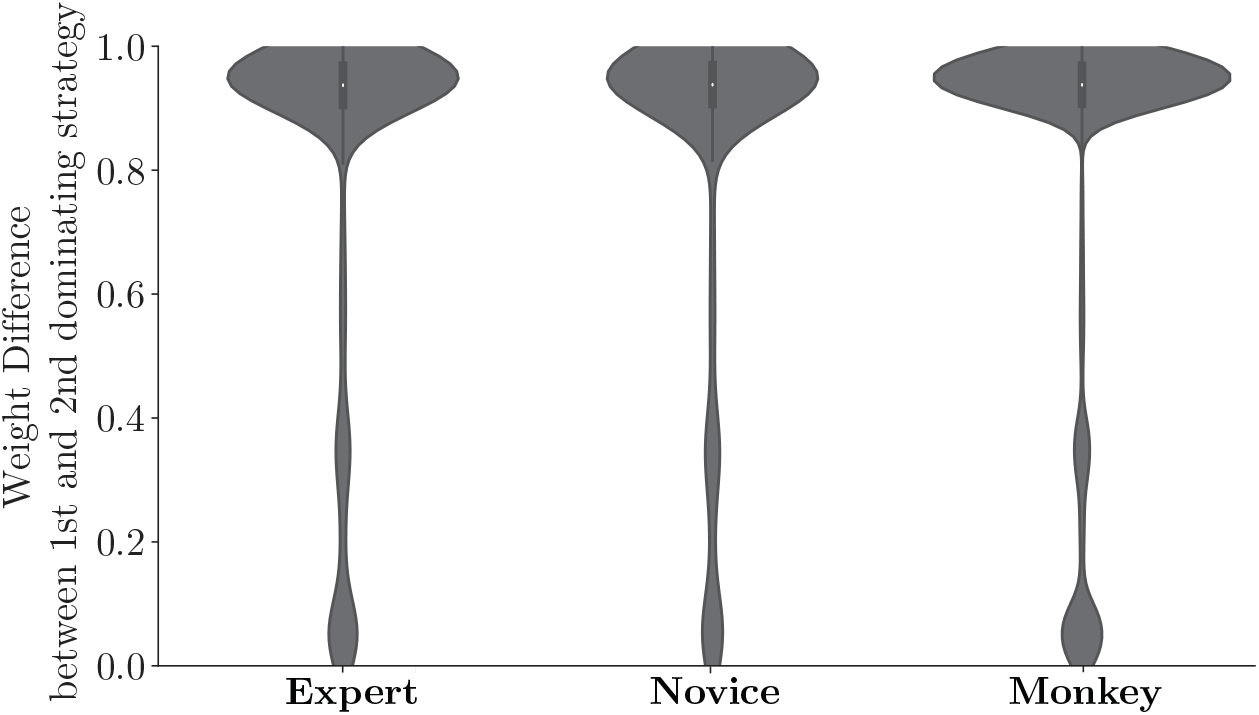
Take-The-Best strategy heuristic. The distribution of the weight difference between the most and the second dominating strategies.

### A.2 LoPS gammar induction

To induce the structure of the Language of Problem-Solving (LoPS) from each subject’s behavioral data, we tackle this inverse problem in three steps: Feature extraction, Strategy fitting, and Grammar induction. The pseudocode can be found in Algorithm 1.

#### A.2.1 Feature extraction

We first extract location information about each game element from the visual input time series. From each frame of the Pac-Man game, we obtain the location of the Pac-Man *l*, the locations of the two ghosts *l*_*g*1_, *l*_*g*2_, the locations of the energizers *l*_*e*_, and the locations of the pellets *l*_*p*_.

We compute the utility values for all directions (𝒜 = left, right, up, down) under each of the seven strategies included in LoPS. Note that not all directions are always available. For unavailable directions, utility values are set to negative infinity. The moving direction is computed according to the largest average utility value for each strategy.

We then determine the utility associated with each direction and its possible trajectories. Let *l* represent Pac-Man’s position and τ(*l, a*) represent a path starting from *l* and moving in the direction of *a* with a length of 10. We compute the utility of path τ(*l, a*) under each strategy as follows (for simplicity, τ(*l, a*) is denoted as τ):

- *local* strategy: 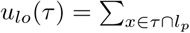 Reward(*x*)
- *energizer* strategy: 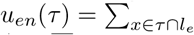 Reward(*x*)
- *save* strategy: 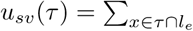 Penalty(*x*)
- *approach* strategy: 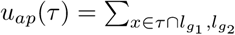 Reward(*x*)
- *evade* strategy: 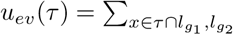 Penalty(*x*), if *g*_1_, *g*_2_ are normal, else 0.

The reward and penalty utilities of each game element in the model can be found in Table C1.

For each direction *a* ∈ 𝒜, its utility *U*_π_(*l, a*) is obtained by averaging all the path utilities *u*_π_(τ) in that direction:

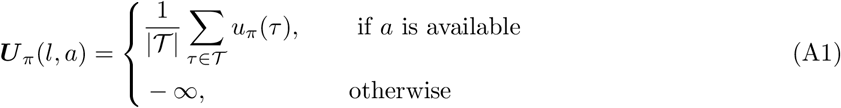

Two strategies, *global* and *stay*, do not depend on utilities associated with Pac-Man location *l*. The utility of *global* strategy *U*_*gl*_ counts the total number of pellets in the entire maze excluding those within five steps of the current position. The utility of *stay* strategy *U*_*st*_ is negative infinite in all directions, leading Pac-Man to stay in place.

Another set of features includes a series of relational-based state variables, capturing the necessary information for strategy initiation and arbitration. These variables include *s*_*e*_ (Dijkstra distance from Pac-Man to the closest energizer), *s*_*g*1_ (Dijkstra distance from Pac-Man to Blinky), and *s*_*g*2_ (Dijkstra distance from Pac-Man to Clyde), *s*_*b*_ (Local pellet number within 5 steps away from Pac-Man), *s*_*m*1_ (Blinky’s mode), and *s*_*m*2_ (Clyde’s mode). To simplify calculations, we binarize the distance and the pellet number variables. Further information about state variables can be found in Table C3.

The pseudocode for feature extraction can be found in Algorithm 2.

#### A.2.2 Strategy fitting

The strategy fitting follows the same procedure described in our previous work [19]. The process of strategy fitting is built upon the assumption that the strategy weights remain stable for a period of time. This period is defined by fine-grained time windows Δ = δ_1_, δ_2_, …, δ_*k*_, separated by essential game events, including changes in Pac-Man’s direction, ghost consumption, energizer consumption, and ghost mode change.

Within each time segment, a softmax policy is employed to linearly combine utilities under each basis strategy, with the strategy weights as model parameters. The probability of choosing a certain action *a* is defined as:

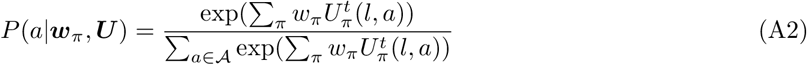

where *w*_π_∈ ℝ^7^ represents the strategy weight.

Given the utility and action time series in each time segment, 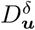, and 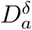, the likelihood function can be formulated as:

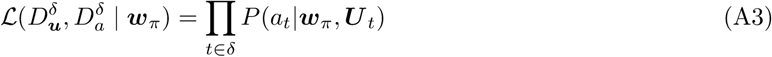

A genetic algorithm (GA) is then used to estimate the weights by maximizing the likelihood function. This implementation of the GA algorithm uses the GA class from the scikit-opt library in Python with the following parameters Population size: 100, mutation probability: 0.1, crossover probability: 0.8, and maximum iteration number: 500.

The strategy with the highest weight is used to create a strategy time series 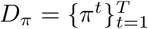, where 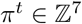 represents the dominant strategy at time *t*. More details about this procedure can be found in Algorithm 3.

The accuracies of joystick movement predictions are based on time points (tiles) and the weights of all strategies.

#### A.2.3 Grammar induction

Given the time series of strategy *D*_π_ and state *D*_*s*_, we induce the UpN-gram G using the structure learning method. The process begins with a grammar graph that only contains uni-grams, 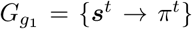. We applied a hybrid network learning algorithm (Session A.6 and Algorithm 8) to infer the uni-gram graph.

With this initial graph 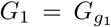, we compute the likelihood score of an alternative graph by concatenating one pair of strategies, *G*_2_. If the likelihood score for the alternative graph is higher than that for the initial graph, we accept the alternative graph as the hypothetical graph. Upon each update of the hypothetical graph, we parse the strategy time series again to generate a new UpN-gram time series *D*_*g*_ based on the updated grammar rules. This new time series is used to compute the likelihood score in the subsequent iteration. This iterative process continues until the likelihood score for the hypothetical graph reaches a maximum. At this point, the induced graph structure is defined as the subject’s LoPS grammar 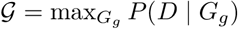. This pseudocode for this procedure can be found in Algorithm 4.

The parsing algorithm used in the grammar induction follows the principle of Minimum Description Length (MDL). Given a specific strategy time series *D*_π_, we parse it into an UpN-gram time series *D*_*g*_ that uses the minimal number of grammar elements according to a set of grammar rules *G*_*g*_. Thereby, we compress the original strategy time series into the shortest possible description. The pseudocode for the Parsing algorithm can be found in Algorithm 5.

#### A.2.4 Validation of LoPS grammar

To validate the LoPS induction algorithm, we construct artificial agents with flexible ground-truth grammar, varying in their grammar complexity (𝒢_1_: uni-grams; 𝒢_2_: bi-grams; 𝒢_3_: tri-grams). For an agent with a defined grammar complexity, their grammars are randomly selected from the library we induce from human and monkey subjects. The artificial agent then interacts with the Pac-Man game environment. We collect *M* = 500 different artificial agents’ gameplay data. We then apply the LoPS induction algorithm (Algorithm 1) to infer each agent’s LoPS grammar. In Supplementary Figure A2, we plotted the accuracy of estimated agent type and grammar complexity, along with the increasing sample size of strategy sequences (*D*_π_). Both estimates can reach high accuracy below the quantity of the experimental data we collected in humans (∼ 4000) and monkeys (∼ 20000).

**Supplementary Figure A2:**
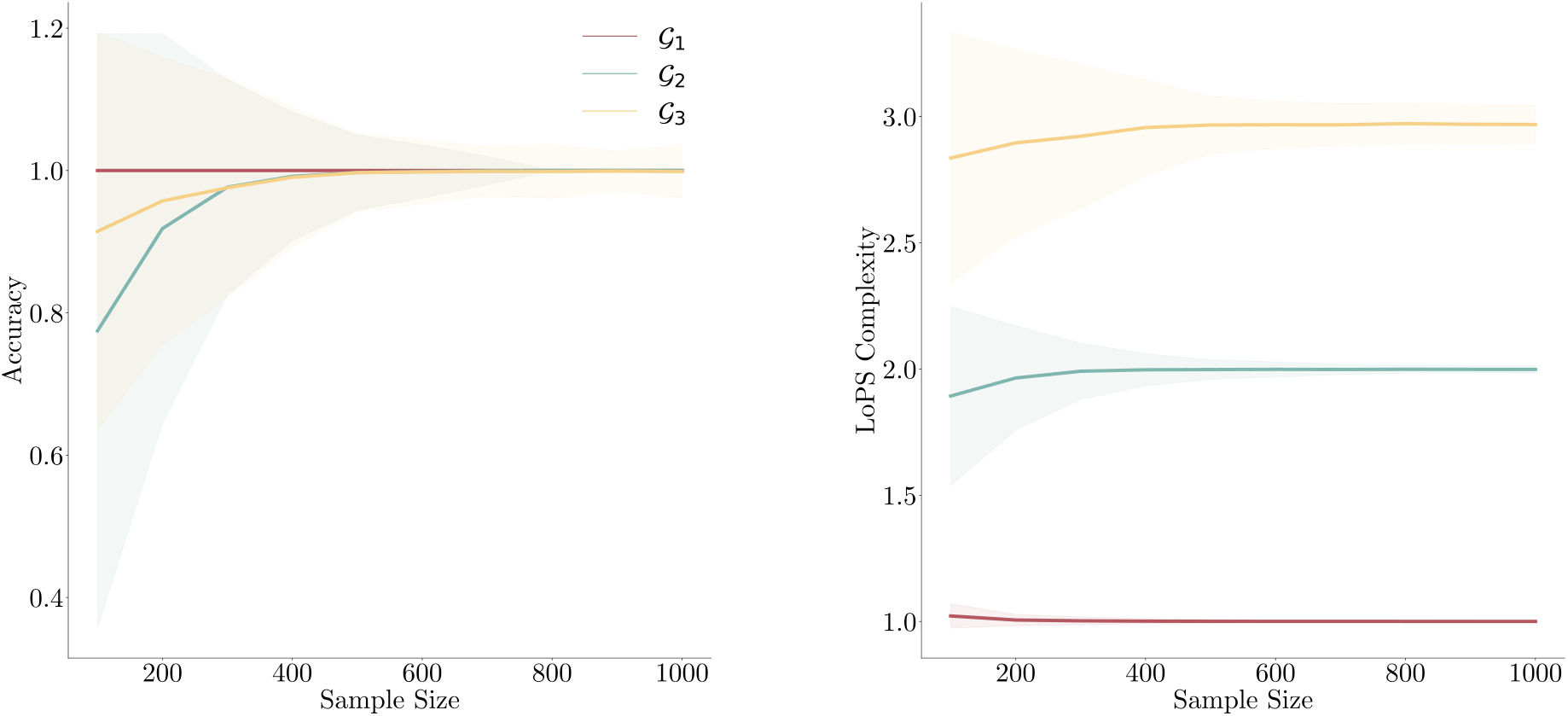
Validation of the LoPS induction algorithm. Three sets of simulated data are generated with different complexity levels, indicated with different colors. The solid line denotes the average estimate. The shade denotes the standard error. The accuracy of estimated agent type (left) and grammar complexity (right) can reach high accuracy below the quantity of the experiment data we collected in humans (∼ 4000) and monkeys (∼ 20000).

### A.3 Structure learning in Probabilistic graphical model

To compute the marginal likelihood of a dataset *D* given a hypothetical graph structure *G* ={𝒱, ℰ},, we integrate over all possible parameter configurations Θ under the given graph structure:

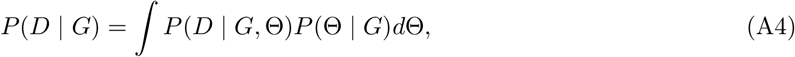

where *P* (*D*| *G*, Θ) is the likelihood of the data given the graph structure and a specific parameter configuration, and *P* (Θ| *G*) is the prior probability of the parameter given the graph structure. The integration over Θ is necessary to avoid favoring more complex models which are characterized by larger parameter sets.

To identify the best network structure given the data, we use a scoring method that seeks to maximize this marginal likelihood over all possible graph structures:

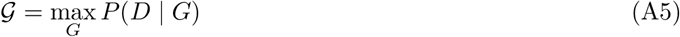

In this setup, 𝒢 denotes the optimal graph structure, which is the one that maximizes the likelihood of the data.

### A.4 PC algorithm for learning a Markov network

For a Markov network with *n* variables 𝒱 = {𝒱 _1_, … 𝒱 _*n*_}, the joint probability is defined as a product of potentials on the subsets of variables (clique) 𝒱 _*c*_ ∈ 𝒱:

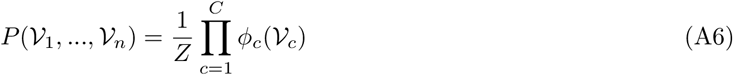

where *Z* is the normalization constant, φ_*c*_ is potential function for the clique 𝒱 _*c*_.

The PC algorithm is a constraint-based structure learning approach [21, 22]. It starts with a fully connected undirected graph and iteratively removes edges based on conditional independence tests. Here’s a brief summary of the method:

1. Start with a fully connected graph.
2. For each pair of variables 𝒱_*k*_ and 𝒱_*l*_, test the conditional independence of 𝒱_*k*_ and 𝒱_*l*_ given the set of all other variables 𝒱_−*k*,−*l*_.
3. If 𝒱_*k*_ and 𝒱_*l*_ are conditionally independent given 𝒱_−*k*,−*l*_, remove the edge between 𝒱_*k*_ and 𝒱_*l*_.
4. Repeat 2-3 until no more edges can be removed, and output the structure of the resulting graph.

The conditional independence test used in the PC algorithm is performed using a Bayesian approach, where the likelihoods under the independence and dependence hypotheses are evaluated with the data. The independence hypothesis assumes that the joint distribution of 𝒱_*k*_ and 𝒱_*l*_ can be factored into separate distributions conditioned on 𝒱_−*k*,−*l*_:

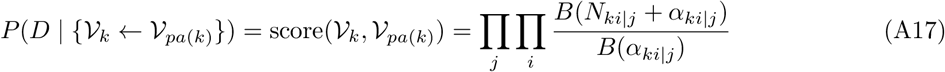

whereas the joint distribution in the dependence hypothesis can’t be factorized:

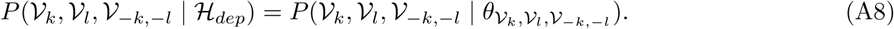

By treating variables as categorical and assuming Dirichlet priors for all parameters, we can compute the marginal likelihood with the dataset 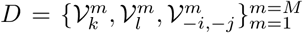 for the independent hypothesis and the dependent hypothesis as

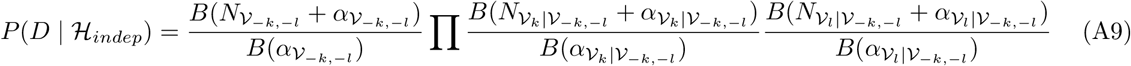

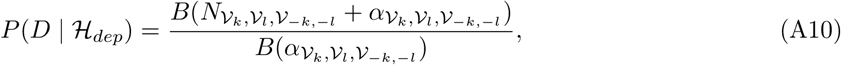

where *B* is beta function, *N*_*x*_ is the number of times that event *x* is presented in the data, α_*x*_ is the corresponding hyperparameter for each event *x* [21, 22].

The pseudocode for the conditional independence test and PC algorithm can be found in Algorithm 10 and Algorithm 6.

To validate the PC algorithm, we first randomly generate Markov graph structures and specify the parameters for their local pairwise potentials. However, it doesn’t mean that the joint distribution fulfills the Markov property that the probability of each variable is dependent only on its immediate neighbors. We compute the following two distributions: *P*(𝒱_*i*_|𝒱_−*i*_) and *P* (𝒱_*i*_|*ne*(𝒱_*i*_)) according to the joint probability of the specified local structures and parameters, and make sure that they are equal for all variables. In this way, we generate *M* Markov networks that fulfill local Markov properties, each characterized by its unique structure and parameters, to serve as the ground truth to validate the PC algorithm.

For each Markov network generated, we create *N* samples under the marginal distributions of each variable, computed according to the defined joint distributions. We then recover the ground-truth network structure by applying the PC algorithm to the synthetic data. Without loss of generality, we set *n* = 3, *M* = 50000, and varying sample size from *N* = 20 to *N* = 2000. The average accuracy of the PC algorithm is depicted in Supplementary Figure A3, where the accuracy is computed according to the congruence between edge sets specified by the ground-truth network and the induced network. The validation results indicate that the PC algorithm may recover the network structure well even with a small sample.

**Supplementary Figure A3:**
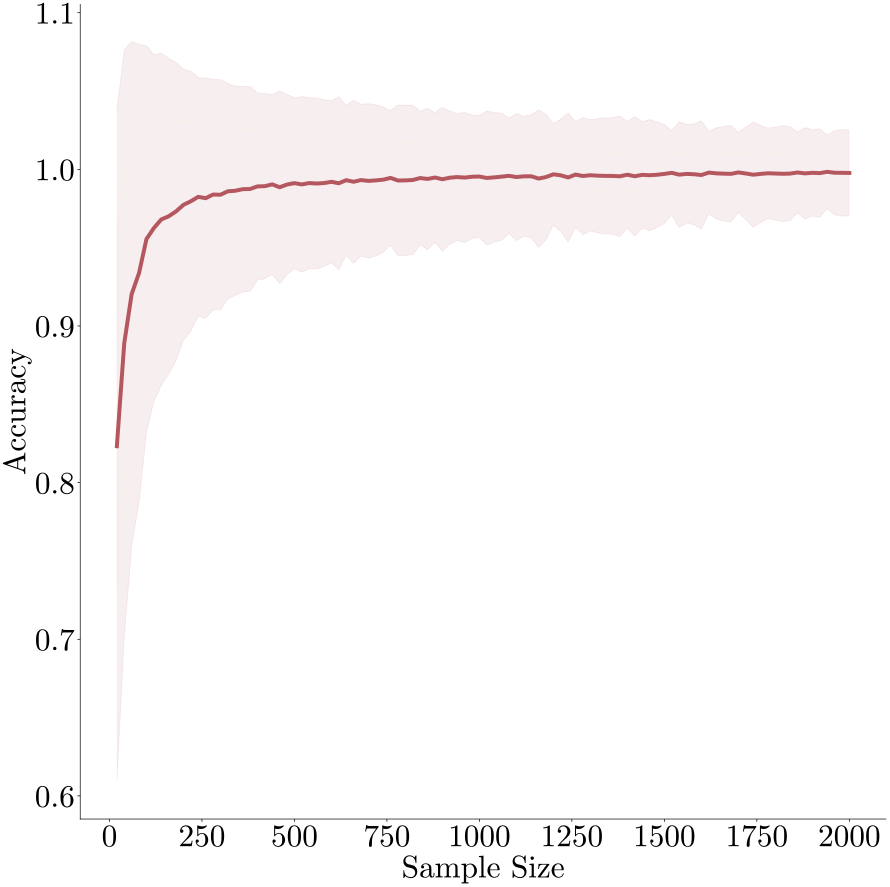
Validation of the PC algorithm on synthetic data generated from the ground-truth Markov network. The solid line denotes the average accuracy. The shade denotes the standard error.

### A.5 Network scoring algorithm for learning a Bayesian network

For a Bayesian network with *n* variables 𝒱 = {𝒱_1_, … 𝒱_*n*_}, the joint probability is defined as

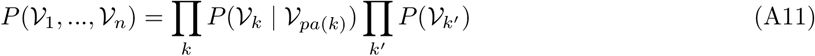

where 𝒱_*pa*(*k*)_ represents the parents of 𝒱_*k*_ in the network, 𝒱_*k*′_ are the most upstream variables in the network who have no parents.

The method for learning the structure of a Bayesian network involves using network scoring techniques. The computation of the marginal likelihood (Eq A4) generally involves an intractable integration. However, by treating variables as categorical and assuming Dirichlet priors for parameters, the marginal likelihood can be analytically calculated [21, 22].

Specifically, these factorized local distributions can be expressed as

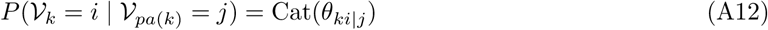

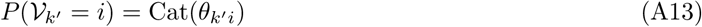

where Cat represents the probability density function of a categorical distribution, θ_*ki*|*j*_ indicates the probability of 𝒱_*k*_ being in state *i* given its parents are in state *j*, and θ_*k*′_ *i* specifies the probability of 𝒱_*k*′_ being in state *i*.

Given data 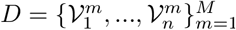, the likelihood of these local distributions can be expressed as:

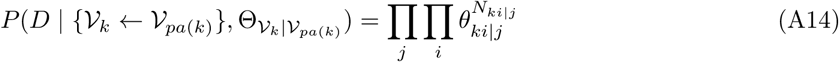

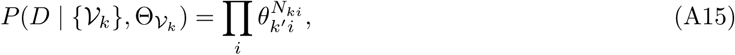

where *N*_*ki*|*j*_ is the number of times that 𝒱_*k*_ is in state *i* and its parents in state *j* in the data, *N*_*k*′_ *i* is the number of times that 𝒱_*k*_ is in state *i*,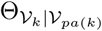 and 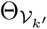 are parameter sets.

We apply the Bayesian Dirichlet likelihood equivalent uniform (BDeu) prior for the parameters:

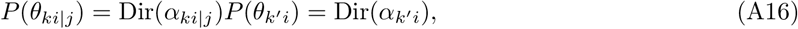

where 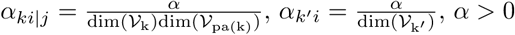. We can compute the marginal local likelihood as

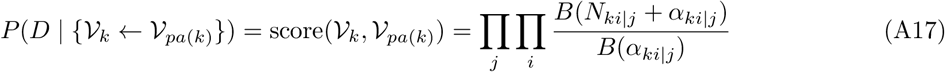

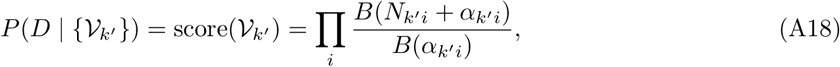

where *B* is the beta function.

Based on these marginal local likelihoods, we can then combine them to compute the likelihood of the whole graph as

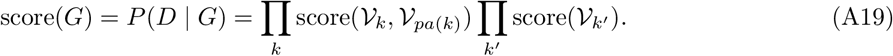

Based on this likelihood score, we can search for graph structures that yield the highest scores. This network scoring approach can be more efficient than the PC algorithm, as it allows for the comparison of differences in local structures between two networks, thanks to the score function’s compositionality into local likelihoods. Therefore, search heuristics that involve the local addition or removal of edges can be particularly effective [22].

The pseudocode for the network scoring algorithm can be found in Algorithm 7.

To validate the network scoring algorithm, we generate Bayesian networks with *n* upstream variables and *m* downstream variables. For each of the downstream nodes, we independently choose one out of the 2^*n*^ possible combinations of the upstream nodes as its parent nodes. We parameterize the conditional probability distribution underlying this local structure as a categorical distribution. Parameters are randomly generated from a uniform distribution whose support is between 0 to 1. In this way, we generate *M* Bayesian networks, each characterized by its unique structure and parameters, to serve as the ground truth for the network scoring algorithm.

For each Bayesian network, we generate *N* samples under defined conditional probability distributions. We then attempt to recover the ground-truth network structure by applying the network scoring algorithm to these synthetic data. Without loss of generality, we set *n* = 3, *m* = 2, *M* = 50000, and varying sample size from *N* = 20 to *N* = 2000. The average accuracy of the network scoring algorithm was depicted in Supplementary Figure A4, where the accuracy is computed according to the congruence between edge sets specified by the ground-truth network and the induced network.

**Supplementary Figure A4:**
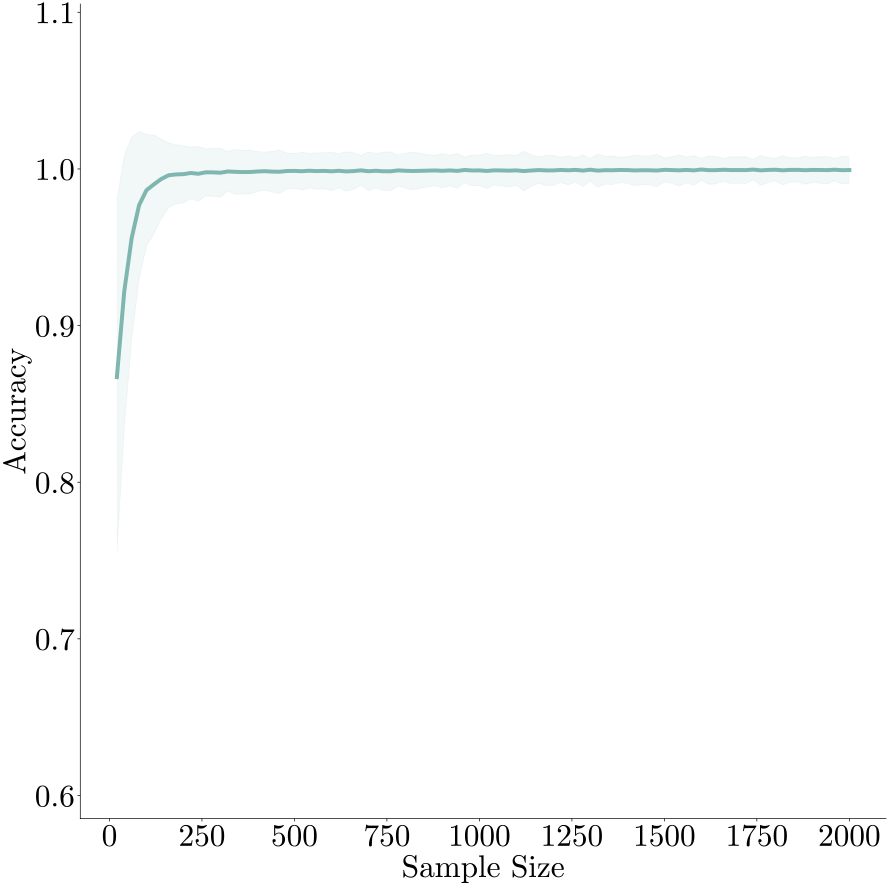
Validation of the network scoring algorithm on synthetic data generated from the ground-truth Bayesian network. The solid line denotes the average accuracy. The shade denotes the standard error.

### A.6 Hybrid network learning

Here, we explore a special case in which upstream variables (states) are modeled through a Markov network, which is an undirected network, while the interconnections between the upstream and the downstream (strategies) variables are captured by a Bayesian network, which is directed. To tackle the challenge of structure learning in such a setting, we have developed a hybrid network learning algorithm, detailed as follows.

First, we apply the PC algorithm to induce the structure of the Markov network (*G*_*m*_) amongst upstream variables. This structure allows the identification of neighboring nodes, *ne* 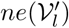, for any given upstream node 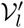. These neighbors form the Markov blanket: the smallest set of nodes that make 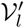 conditionally independent to all other nodes.

Subsequently, we apply the network scoring algorithm to induce the structure of the Bayesian network (*G*_*b*_). The computation of the marginal likelihood entails a directed edge from upstream node 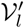 to a downstream node 𝒱_*k*_, necessitates the additional conditioning on the neighboring nodes *ne*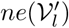:

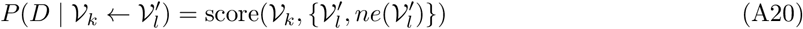

This score function is computed according to Eq A17. Consequently, the revised likelihood score function is:

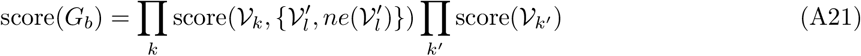

With this likelihood score, we search for the graph structure that yields the highest score.

The hybrid network learning algorithm exploits the conditional independence properties of the Markov network to reduce the computational complexity, transitioning from conditioning on all upstream state variables to conditioning solely on the Markov blanket.

The pseudocode for the hybrid network learning algorithm can be found in Algorithm 8.

To validate the hybrid network learning algorithm, we generate synthetic data similar to the scenario in the Bayesian network validation. However, here, the upstream variables are not independent. Instead, they are generated from the Markov network defined in Session A.4. Without loss of generality, we set *n* = 3, *m* = 2, *M* = 50000, and varying sample size from *N* = 20 to *N* = 2000. The average accuracy of the hybrid network learning algorithm is depicted in Supplementary Figure A5, where the accuracy is computed according to the congruence between edge sets specified by the ground-truth network and the induced network. Our hybrid network learning algorithm can accurately capture the true network structure used to create the simulated data.

**Supplementary Figure A5:**
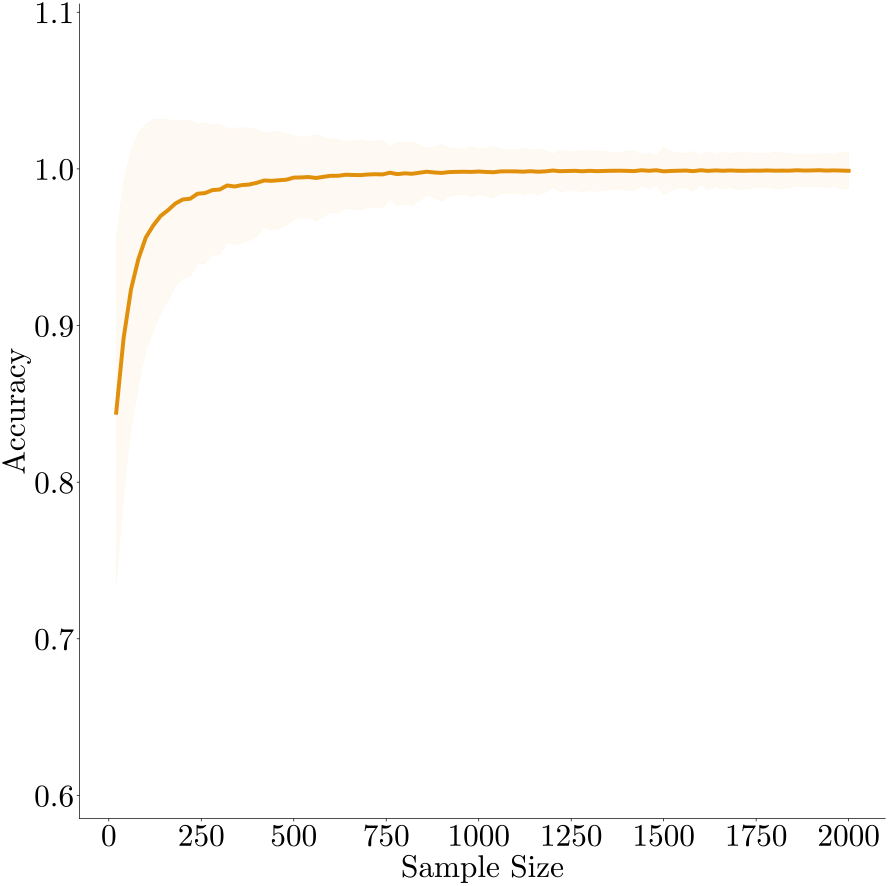
Validation of the hybrid network learning algorithm on synthetic data generated from the ground-truth hybrid network. The solid line denotes the average accuracy. The shade denotes the standard error.

## Appendix B Videos

Video 1: example game video from example human 1 (novice).

Video 2: example game video from example human 2 (expert).

Video 3: example game video from Monkey P.

Video 4: example game video for a skip-gram grammar from example human 2

Video 5: example game video video for a tri-gram *Stay-Energizer-Approach* from example human 2.

## Appendix C Tables

**Table C1:** Rewards and penalties in the Pac-Man game.

| Reward (pellet) | Reward (energizer) | Reward (scared ghost) | Penalty (death) | Penalty (save energizer) |
| --- | --- | --- | --- | --- |
| 2 | 4 | 10 | -10 | -4 |

**Table C2:** Concepts in Language of Problem-Solving.

|  |  |
| --- | --- |
| $I$ | Visual inputs |
| $a$ | Joystick movement actions, $a = \{\text{up, down, left, right}\}$ |
| $U$ | Utilities associated with each action under a specific strategy |
| $s$ | Game states, $s = \{s_e, s_b, s_{g1}, s_{g2}, s_{m1}, s_{m2}\}$, categorical variable denoting the condition of the current game states. Detailed definitions can be found in Table C3 |
| $\pi$ | Strategy, selected from a set of seven strategies. Detailed definitions can be found in Table C4 |
| $\mathcal{G}$ | Grammar, specifies the temporal dependency between the strategies. $\{g_1, g_2, g_3, \dots, g_{skip}\} \in \mathcal{G}$, where $g_1$ is the uni-gram, $g_2$ is the bi-gram, $g_3$ is the tri-gram, etc. $g_{skip}$ is the skip-gram |
| $C_{\mathcal{G}}$ | LoPS Complexity, defined as average n-gram length, $C_{\mathcal{G}} = \langle n \rangle_{P(g_n)}$, where $P(g_n)$ is the probability of each n-gram based on the parsed n-gram sequence according to the subjects' grammar $\mathcal{G}$ |

**Table C3:**
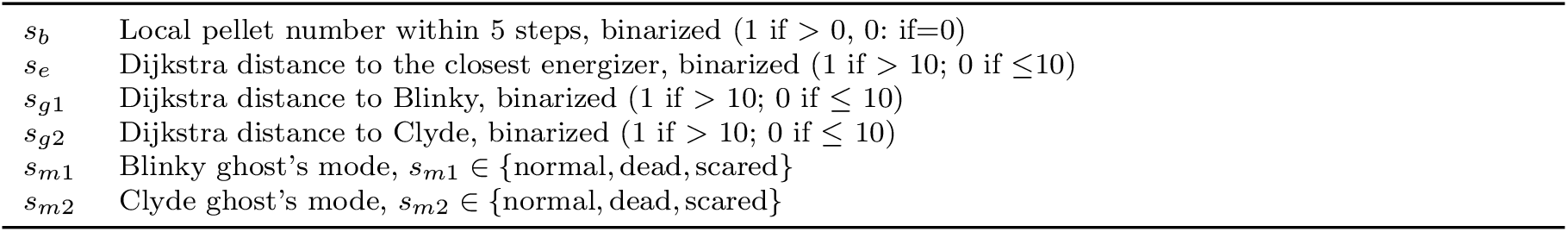
Game state variables.

**Table C4:** Strategies.

|  |  |
| --- | --- |
| $\pi_{lo}$ | *local* strategy moves Pac-Man in the direction that offers the largest reward within a 10-tile radius |
| $\pi_{gl}$ | *global* strategy moves Pac-Man in the direction that has the largest total number of pellets in the whole maze |
| $\pi_{en}$ | *energizer* strategy moves Pac-Man toward the closest energizer |
| $\pi_{ap}$ | *approach* strategy moves Pac-Man toward the ghosts, regardless of the ghosts mode |
| $\pi_{ev}$ | *evade* strategy directs Pac-Man away from the nearest ghost. |
| $\pi_{st}$ | *stay* strategy keeps Pac-Man near a specific location |
| $\pi_{sv}$ | *save* strategy keeps Pac-Man away from the energizer |

**Table C5:** Probabilistic Graphical Model.

|  |  |
| --- | --- |
| $G$ | a graph including nodes and edges, $G = \{\mathcal{V}, \mathcal{E}\}$. |
| $\mathcal{V}$ | Nodes in a graph, indexed by $k, l$. The states of a node is indexed by $i, j$ |
| $\mathcal{E}$ | Edges in a graph, categorized into directed ($\mathcal{V}_k \rightarrow \mathcal{V}_l$) or undirected ($\mathcal{V}_k - \mathcal{V}_l$) |
| $\mathcal{V}_{-k, -l}$ | the subset of variable nodes except for $\mathcal{V}_k$ and $\mathcal{V}_l$ |
| $pa, ch$ | Parent, child nodes in a Bayesian network |
| $nb$ | Neighboring nodes in a Markov network |
| $\Theta$ | Parameter set for a graph |
| $D$ | Data, $D = \{\mathcal{V}_1^m, \dots, \mathcal{V}_K^m\}_{m=1}^M$ contains $M$ samples of $K$ variables in a graph |
| $N_x$ | the number of times that some event $x$ is presented in the data |
| $\alpha_x$ | hyperparameter for each event $x$ when using the Dirichlet priors |

## Appendix D Psudocodes for algorithms

### Algorithm 1

LoPS induction algorithm

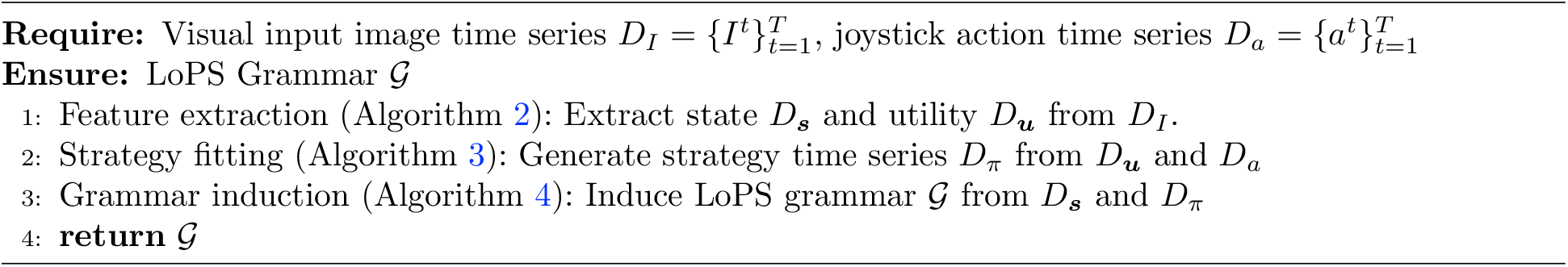

### Algorithm 2

Feature extraction algorithm

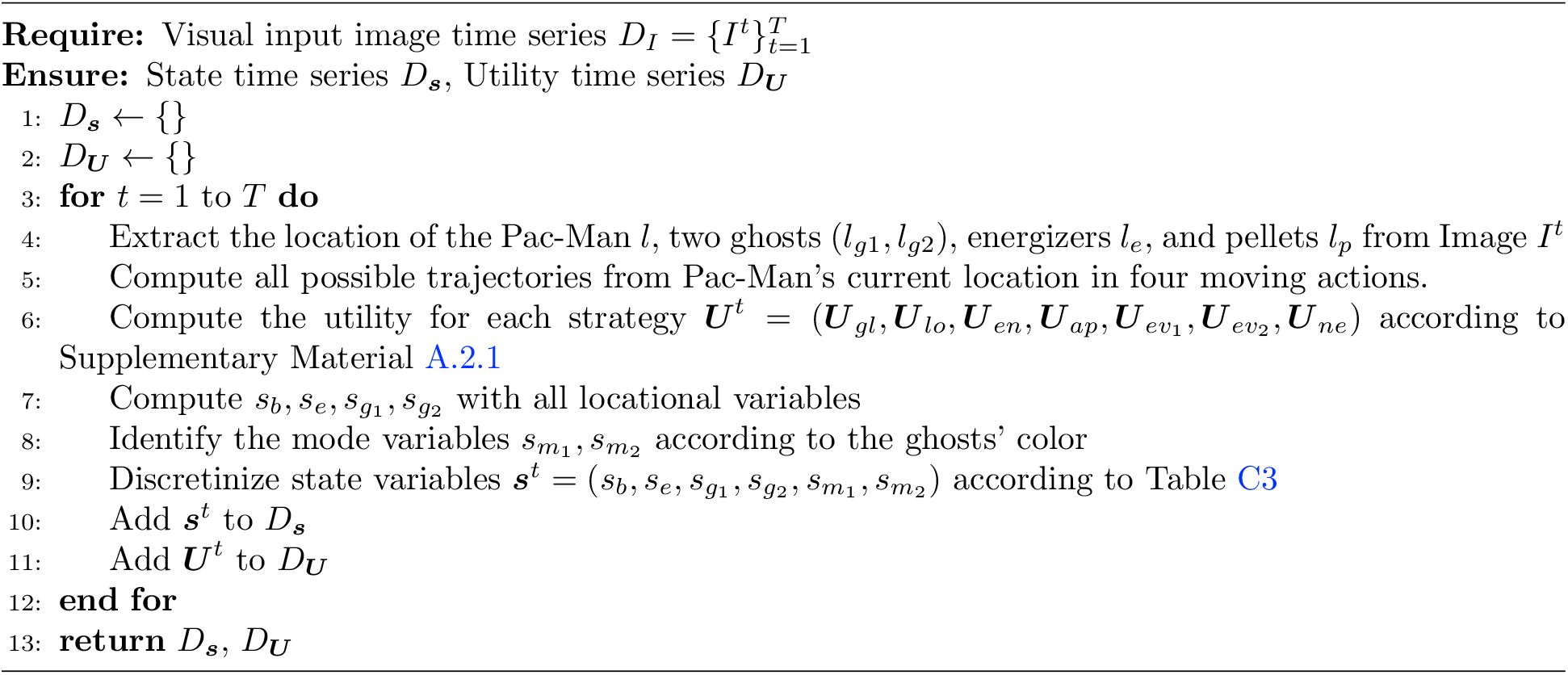

### Algorithm 3

Strategy fitting algorithm

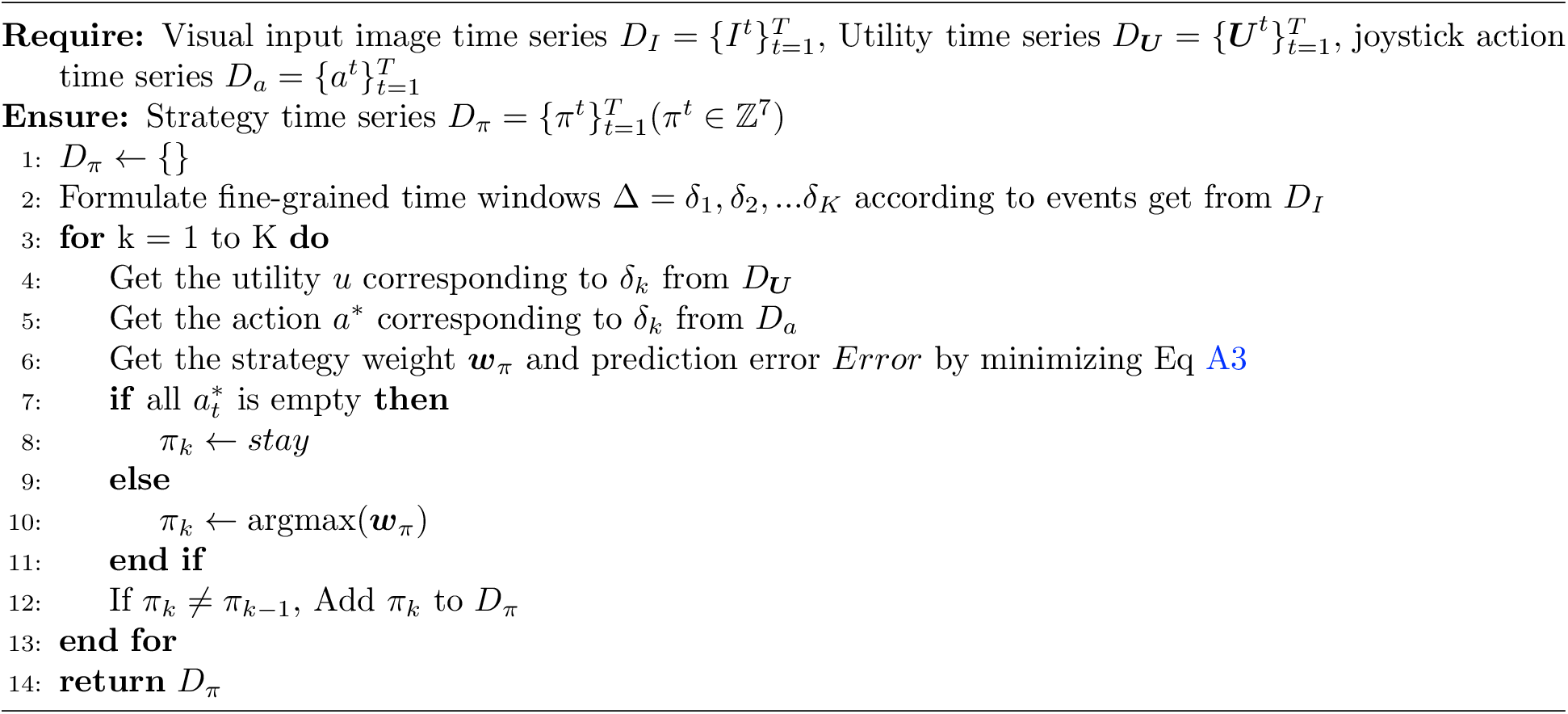

### Algorithm 4

Grammar induction algorithm

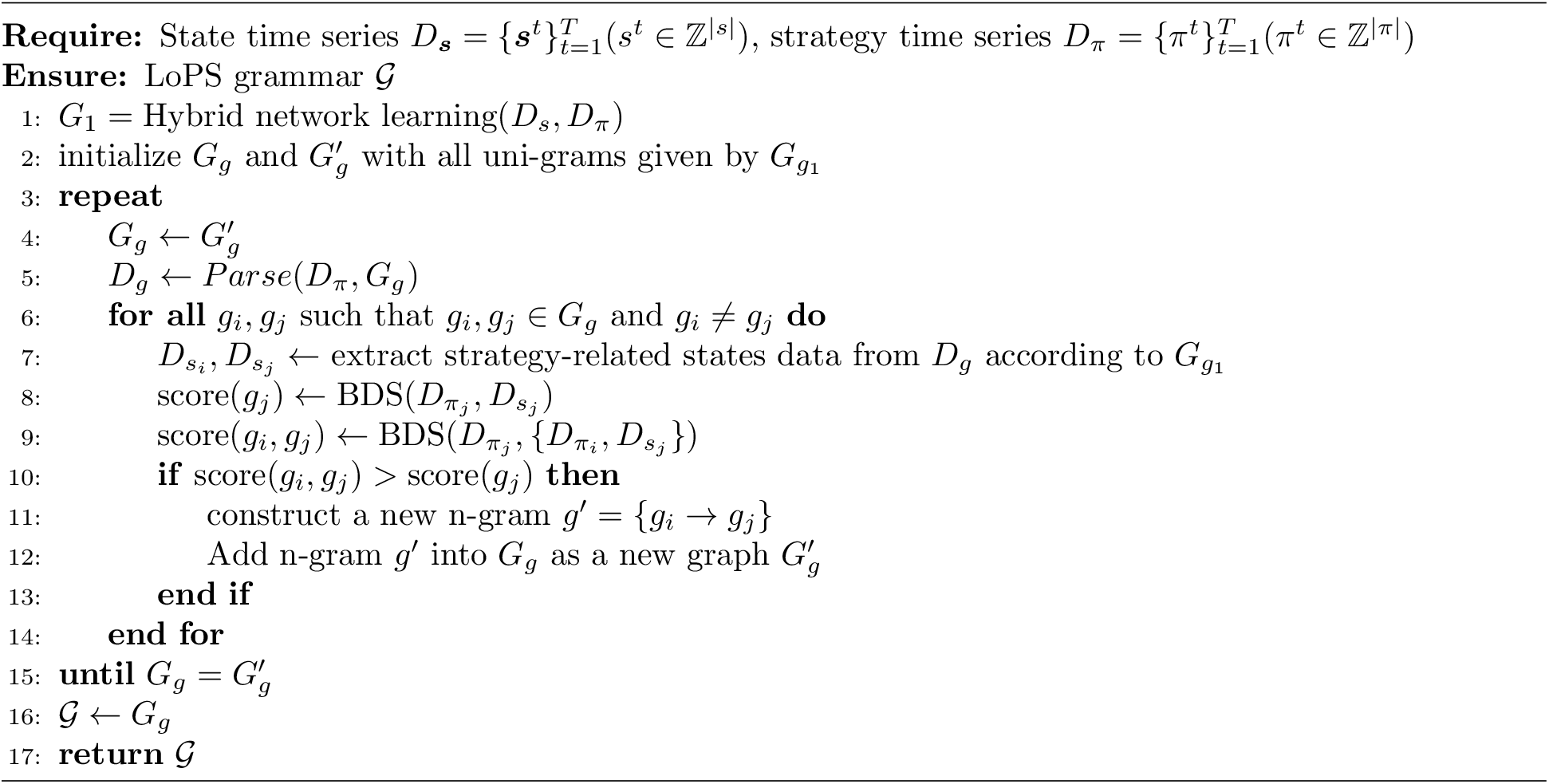

### Algorithm 5

Parsing algorithm *Parse*

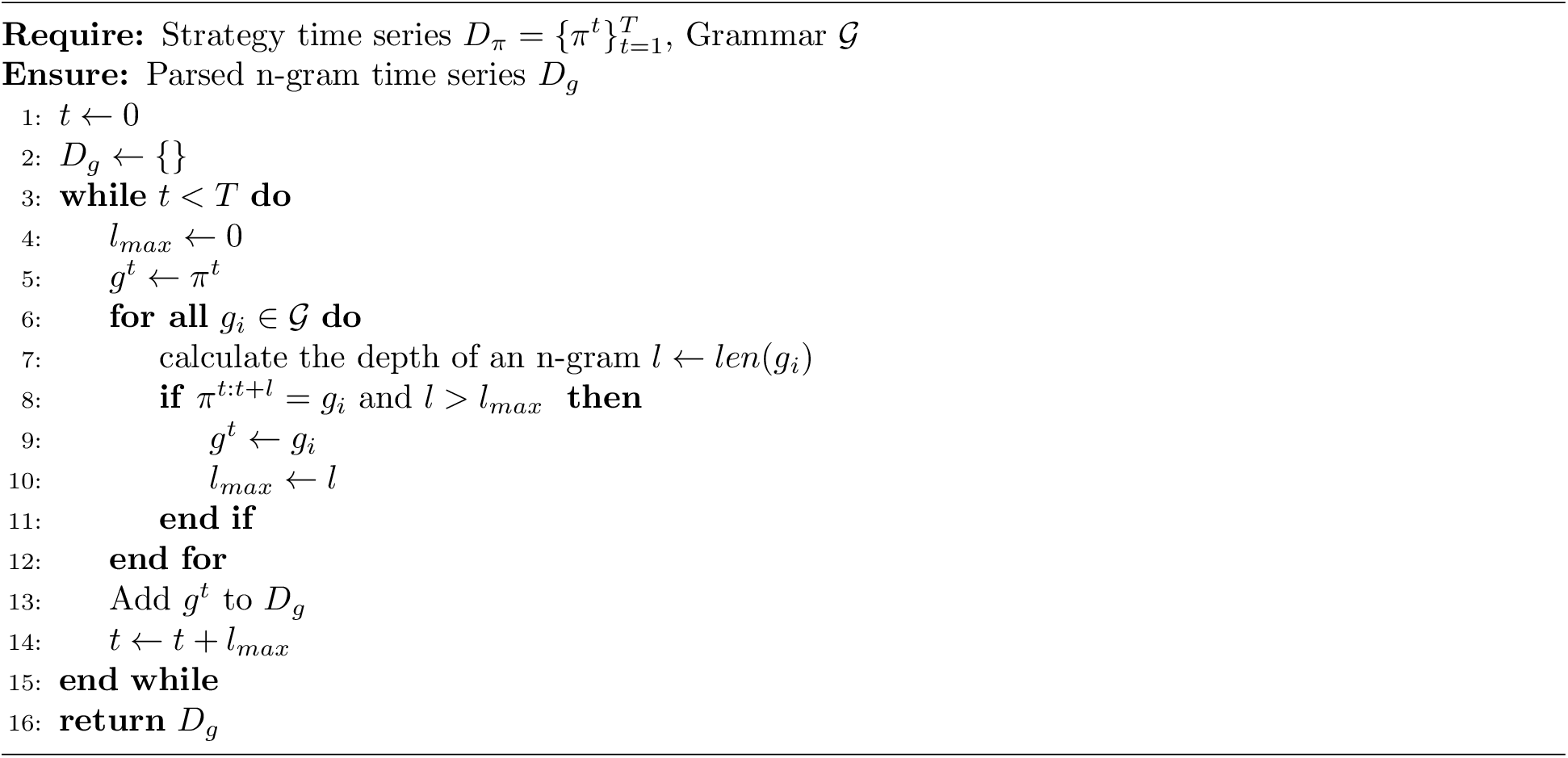

### Algorithm 6

PC Algorithm

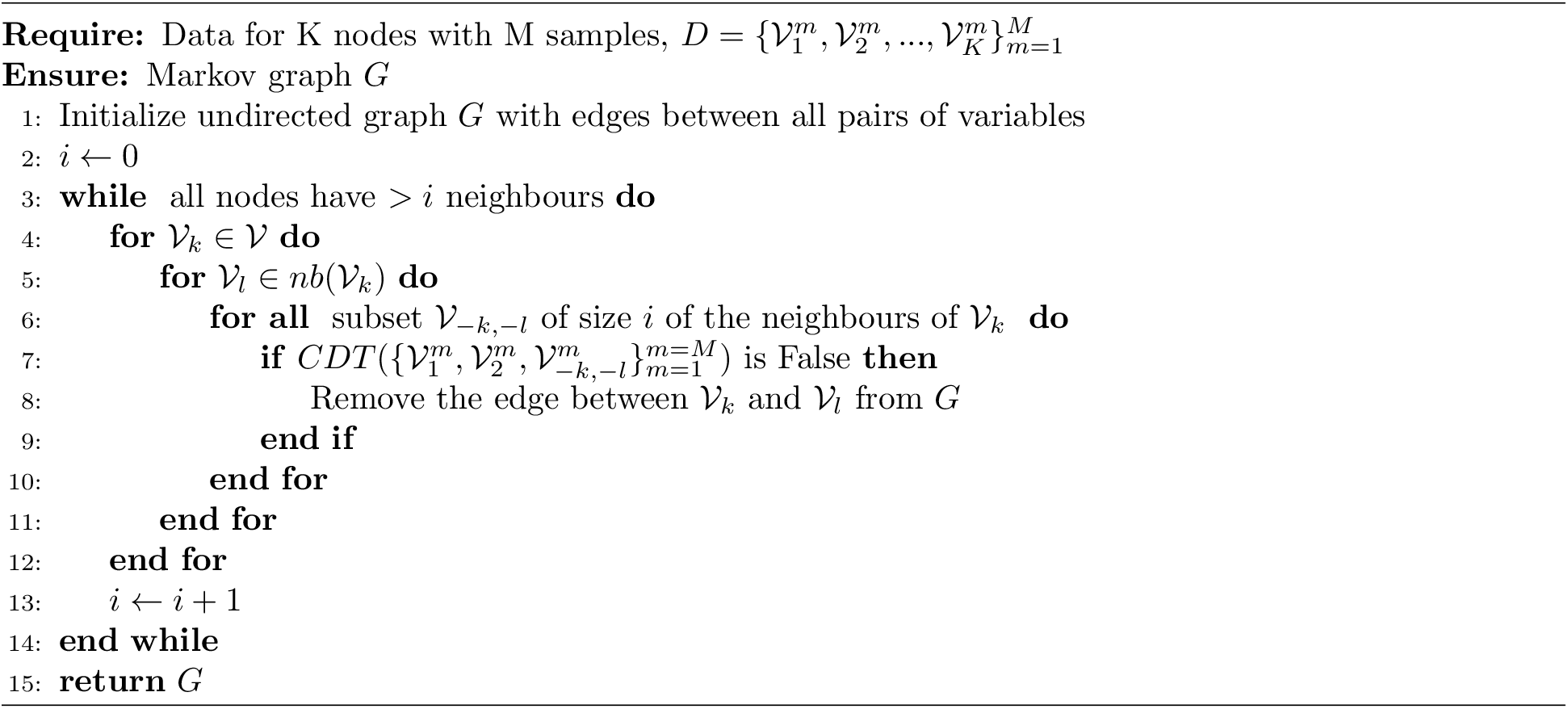

### Algorithm 7

Bayesian network learning

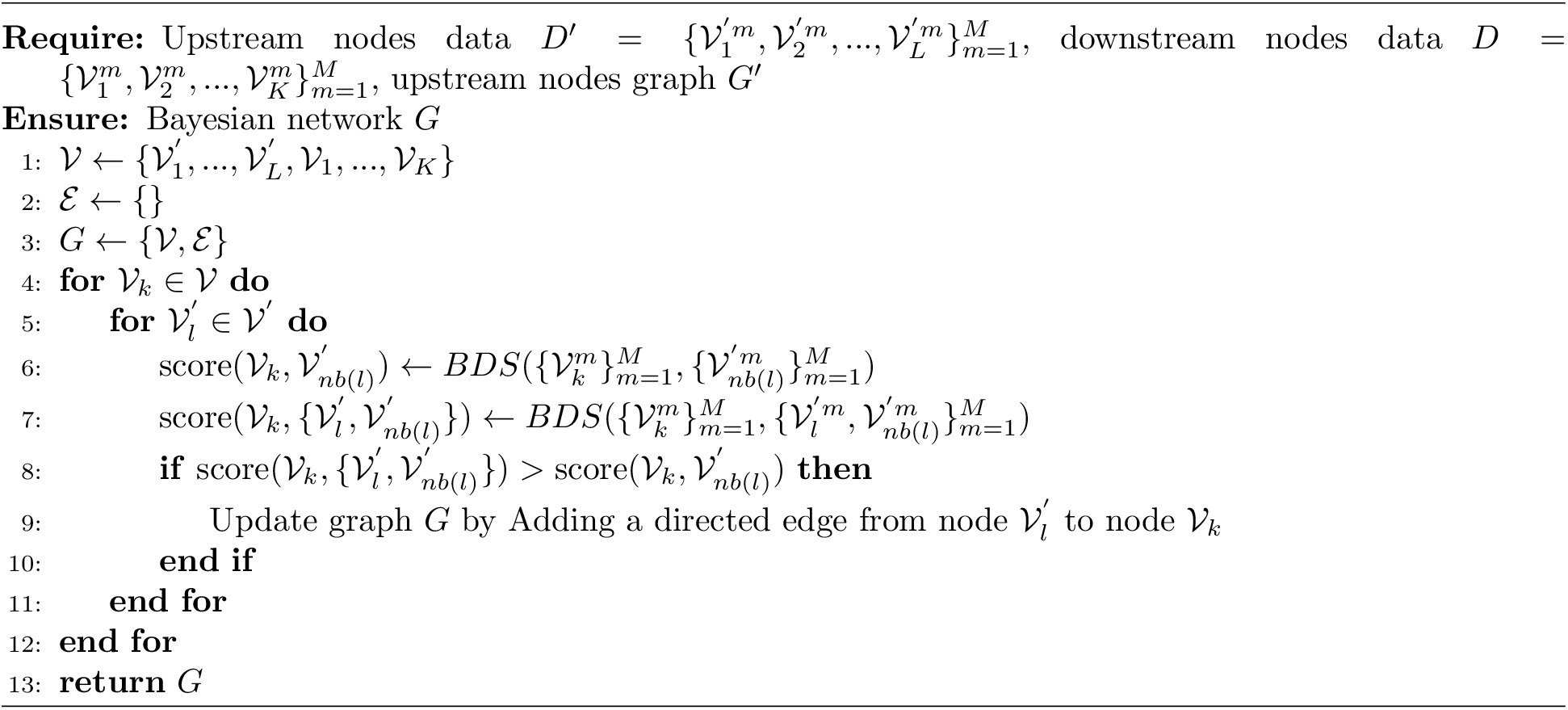

### Algorithm 8

Hybrid network learning

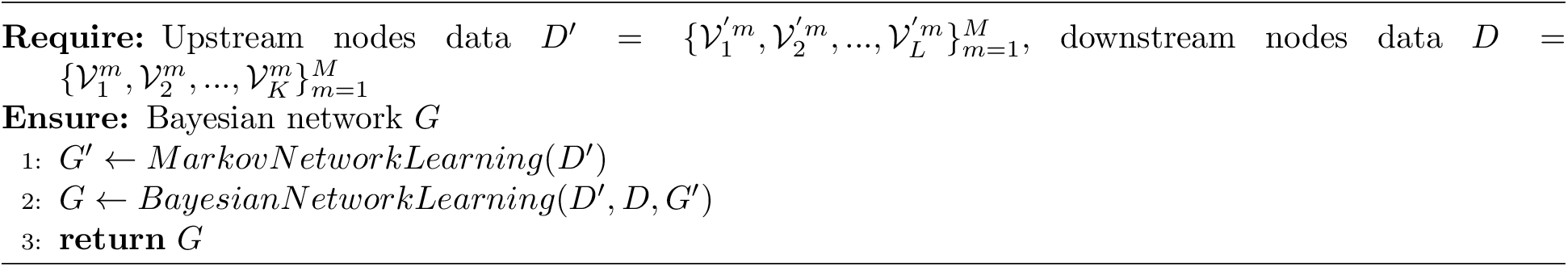

### Algorithm 9

Bayesian Dirichlet Score *BDS*

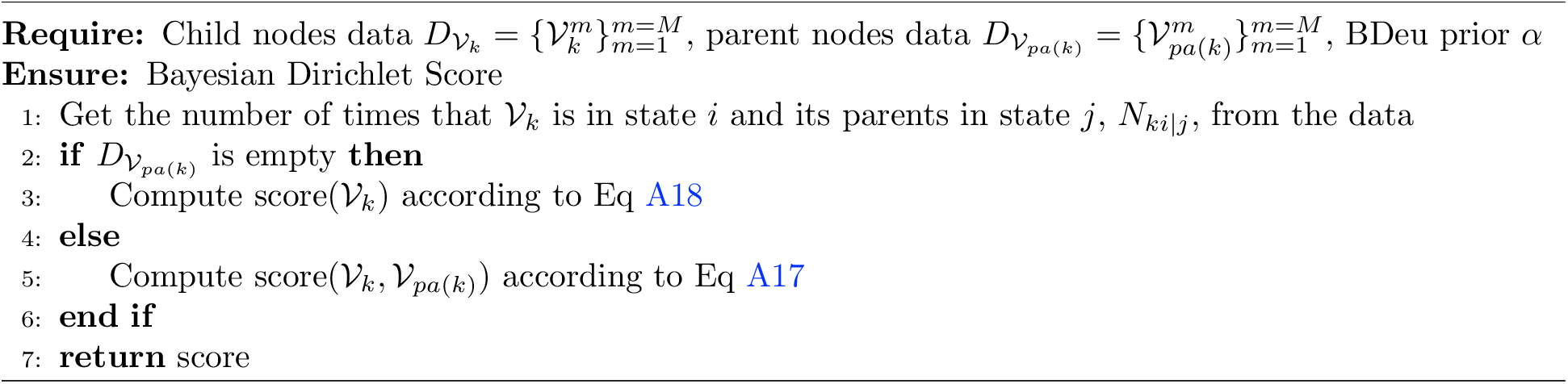

### Algorithm 10

Conditional dependence test *CDT*

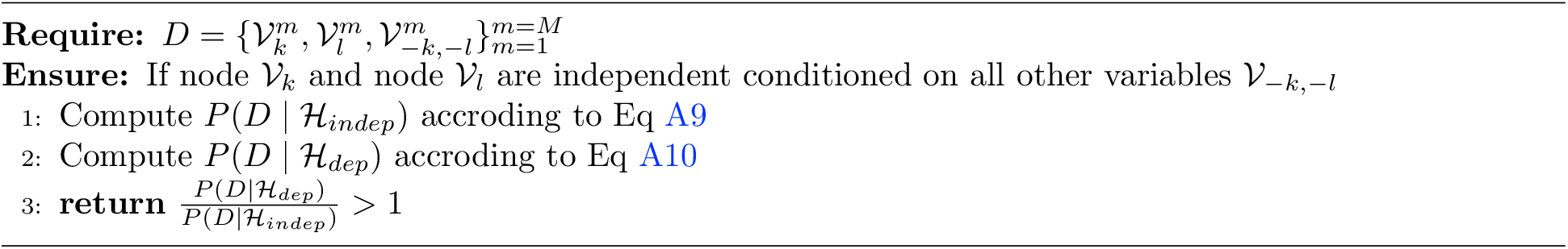

## Notes

https://github.com/superr90/LoPS

### Competing Interest Statement

The authors have declared no competing interest.

